# Biodesign of Frugal CRISPR Kits for Equitable and Accessible Education

**DOI:** 10.1101/2023.10.14.562372

**Authors:** Marvin C. Collins, Matthew B. Lau, William Ma, Aidan Shen, Marie La Russa, Lei S. Qi

## Abstract

Equitable and accessible education on life sciences and bioengineering is crucial to training the next generation of scientists, fostering transparency in public decision-making, and ensuring biotechnology democratization that can benefit a wide-ranging population. As a ground-breaking technology for genome engineering, CRISPR has transformed research and therapeutics. However, hands-on exposure to this technology for educational purposes has remained restricted, primarily due to extensive resources required to execute CRISPR experiments. In this study, we develop an accessible and frugal CRISPR kit, tailored for K-12 education settings. Our CRISPR kit eliminates the need for equipment, prioritizes biosafety, and utilizes cost-effective reagents. By combining CRISPRi gene regulation, chromoproteins, cell-free transcription-translation systems, and smartphone-based quantification, our kit offers a user-friendly approach and a reliable assessment of CRISPR activity, eliminating the need for a traditional laboratory setup. Experiments conducted by high school students in real-world settings highlight the kit’s utility for conducting reliable CRISPR experiments. The frugal CRISPR kit provides a modular and expandable platform to offer hands-on experience in genome engineering, and will facilitate equitable and accessible education and technology democratization for communities of diverse socioeconomic and geographic backgrounds.

## Introduction

Tools and resources to facilitate accessible K-12 (from kindergarten to high school) education in life sciences and bioengineering are crucial for fostering a more informed and equitable society^1-3^. Via accessible education, we not only inspire the next generation of scientists but also create citizens who can understand the complex language behind cutting-edge biotechnologies, thus promoting transparency and responsibility in public decision making while mitigating confusion and apprehension. This process also promotes biotechnology democratization and ensures that advances in life sciences and bioengineering not only benefit a select few, but for everyone regardless of their socioeconomic background, including individuals with low technical and financial resources.

Despite the benefits of promoting accessible K-12 education, major barriers remain in promoting accessible education related to cutting-edge biotechnologies. This is largely due to the complex nature of these technologies, the need for expensive resources, and the biosafety concerns associated with hands-on experience with these technologies. These challenges are further compounded by socioeconomic and geographic disparities in access to quality educational resources. As a result, education on bioengineering in the K-12 setting has been largely confined to theoretical content presented in classrooms.

CRISPR-Cas gene editing technology has recently revolutionized biomedical research and therapeutics^4–6^. Researchers have shown that CRISPR-based gene therapy can restore function of the oxygen-carrying hemoglobin gene, leading to a potential permanent cure of sickle cell anemia^7,8^. Beyond clinical applications, in the laboratory setting, this technology has allowed researchers to alter the genetic code of various organisms, including animals, plants, and microbes, opening a door to advanced genetic engineering that would have been impossible a decade ago^9,10^. Despite its monumental impact on biological sciences at a university and research institute level, hands-on experience with the technology has been absent from high-school curricula, due to resource requirements for carrying out CRISPR experiments (**Fig. 1a**). For example, in the current CRISPR experiment setup, researchers need to invest in expensive laboratory equipment, often costing tens of thousands of US dollars. Furthermore, they need to address biosafety requirements for dealing with cells and biohazards and follow a relatively complex experimental protocol. These barriers are insurmountable to most high schools, leaving most K-12 students with minimal hands-on experiences with CRISPR technology until university.

**Figure 1:**
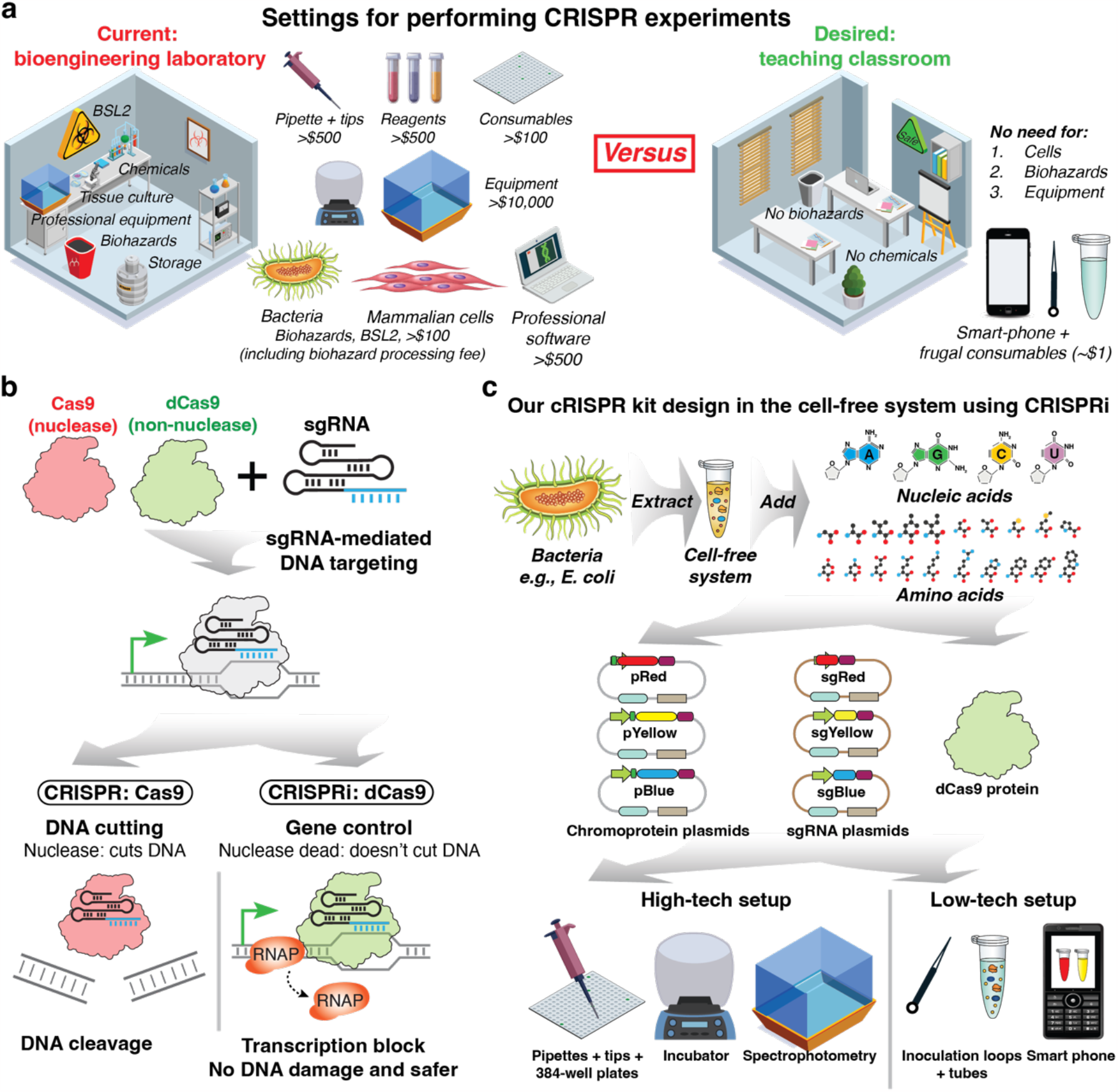
The concept and design of accessible CRISPR kits for K-12 education. **a**. The need for frugal CRISPR kits for accessible education with minimal resource requirements. A comparison between two setups for performing CRISPR experiments. Settings on the left show a biosafety level (BSL) 2 laboratory that is usually required to carry out genome editing experiments, involving various chemicals, tissue culture, storage, biohazards, and professional equipment such as incubator and fluorescence measurement equipment. To perform a CRISPR experiment, students need to get access to pipettes and pipette tips (>$500), various reagents for molecular cloning and tissue culture (>$500), consumables such as 384-well plates (>$100), cells such as bacteria or mammalian cells, biohazards often in the BSL2 setting, equipment (>$10,000), and software for data analysis (>$500). These resources are not easily available to most schools and students. The setting on the right shows an ‘ideal’ frugal setting, involving a routine teaching classroom. No cells, biohazards, or professional equipment is needed. **b**. Illustration of mechanisms of Cas9-mediated genome editing (red) and nuclease-dead dCas9-mediated (green) genome repression (termed CRISPR interference or CRISPRi). Both molecules can bind to specific DNA targets via single guide RNA (sgRNA)-mediated RNA-DNA complementarity, with a nearby protospacer adjacent motif (PAM). Upon binding to DNA targets, Cas9 cuts DNA into two fragments. On the contrary, dCas9 acts as a transcription block to sterically hinder the binding or progression of RNA polymerase (RNAP) on the target DNA, thus repressing transcription of the target gene encoded by DNA, without DNA cleavage. **c**. Our CRISPR kit design is based on CRISPRi that represses gene expression outcomes of the target DNA. Briefly, bacterial extract was supplemented with nucleic acids and amino acids to prepare an in vitro cell-free system (CFS), which was mixed with in-house made plasmids that encode chromoproteins with various colors (pRed, pYellow, pBlue), plasmids that encode chromoprotein-specific sgRNAs (sgRed, sgYellow, sgBlue), and purified dCas9 protein. We compare two experimental setups, one is termed high-tech involving traditional laboratory apparatus and equipment such as pipettes, 384-well plates, incubator, and fluorescence spectrophotometry, and another is termed low-tech involving a smartphone, inoculation loops, and tubes.

While previous studies have successfully reduced the requirements for performing a CRISPR experiment by using a cell-free system, they still require equipment (e.g., an incubator or imager), expensive apparatus (e.g., pipettes), and specialized software (e.g., data analysis)^11–14^. From the perspective of accessibility and sustainability, we reasoned that eliminating these requirements is key to offering hands-on CRISPR experiences to high school students in a more equitable setting (**Fig. 1a**). It is therefore our goal to develop a frugal CRISPR kit that eliminates the need for equipment, minimizes biosafety concerns, and utilizes affordable materials. In this study, we present a safe, frugal, accessible CRISPR kit with no equipment requirement to offer hands-on experiences on CRISPR experiments. Since many high school students have access to smartphones, we intend to measure and quantify the CRISPR activity using smartphones. It is scalable to most high school curricula in a low-resource setting and facilitates accessible education on cutting-edge gene-editing technology.

## Results

### The design of a safe, frugal, and accessible CRISPR kit for education

We analyzed two potential mechanisms for implementing CRISPR experiments in the educational setting (**Fig. 1b**). One mechanism is gene editing that uses the nuclease Cas9 for DNA cleavage^15–17^, and the other is gene regulation that uses the nuclease-dead dCas9 for controlling transcription of the target gene, a method termed CRISPR interference (CRISPRi)^18^. Both mechanisms are based on single guide RNA (sgRNA)-directed specific DNA targeting and can generate visualizable outcomes on the genes encoded by the target DNA. However, since Cas9 is an active enzyme that may have more profound impacts on the environment due to its ability to permanently alter DNA sequences, we proposed to focus on dCas9, which only transiently affects gene expression.

The traditional method of performing CRISPR experiments necessitates the execution of many costly and time-consuming steps, including transforming competent cells, plating transformed cells, picking colonies, and growing the cells overnight. Our proposed CRISPR kit utilizes a transcription-translation-based method to generate an in vitro experimental environment, termed cell-free system (CFS), which has been employed for measuring gene expression and CRISPR gene editing^11,19-21^ (**Fig. 1c**). By using CFS reagents in place of living cells, we eliminate the biosafety and logistical concerns of live-cell transportation and culture.

We further linked *Streptococcus pyogenes* dCas9-based CRISPRi activity to a pigmentation readout via the control of chromoproteins, which are pigmented proteins that do not require special light wavelengths or filters to visualize. We chose three chromoproteins with visually distinct colors that resemble the RGB color mode (red, yellow/green, and blue): eforRed^22^, fwYellow^23^, and aeBlue^24^. We designed single guide RNAs (sgRNAs) that specifically target the protein-coding sequence of each chromoprotein to achieve efficient and specific transcriptional repression.

Using the cell-free system, in the absence of dCas9 or an sgRNA, the transcription and translation machinery will produce chromoproteins that appear as visible pigment. When dCas9 and targeting sgRNA are present, they form a ribonucleoprotein complex to repress transcription of the chromoprotein, reducing the intensity of pigmentation. We measure activity of the CRISPR kit by quantifying the level of pigmentation change. We also compared two setups: one is termed the ‘high-tech’ setup and uses pipettes (for liquid transfer), an incubator (for precise temperature control), and a fluorescence plate reader (for measurement); the other is termed the ‘low-tech’ setup and uses inoculation loops (for liquid transfer), no temperature control, and a smartphone (for measurement) (see **Methods**).

### Testing the performance of the single-color CRISPR kit in the laboratory setup

We first designed specific sgRNAs for each chromoprotein. There are two options for designing an sgRNA targeting a double-stranded DNA: targeting either the template or non-template strand. Previous studies have shown that only non-template-strand-targeting sgRNA can repress transcription, due to steric hindrance between dCas9 and the RNA polymerase^18,25^ (**Fig. 2a**). We therefore designed sgRNAs that target the non-template strand of each chromoprotein, using the CRISPR-ERA algorithm to select sgRNAs without off-target binding sites. We encoded each chromoprotein and each sgRNA under the control of a strong s70 promoter (J23119) (**Fig. 2b**, see **Methods**).

**Figure 2:**
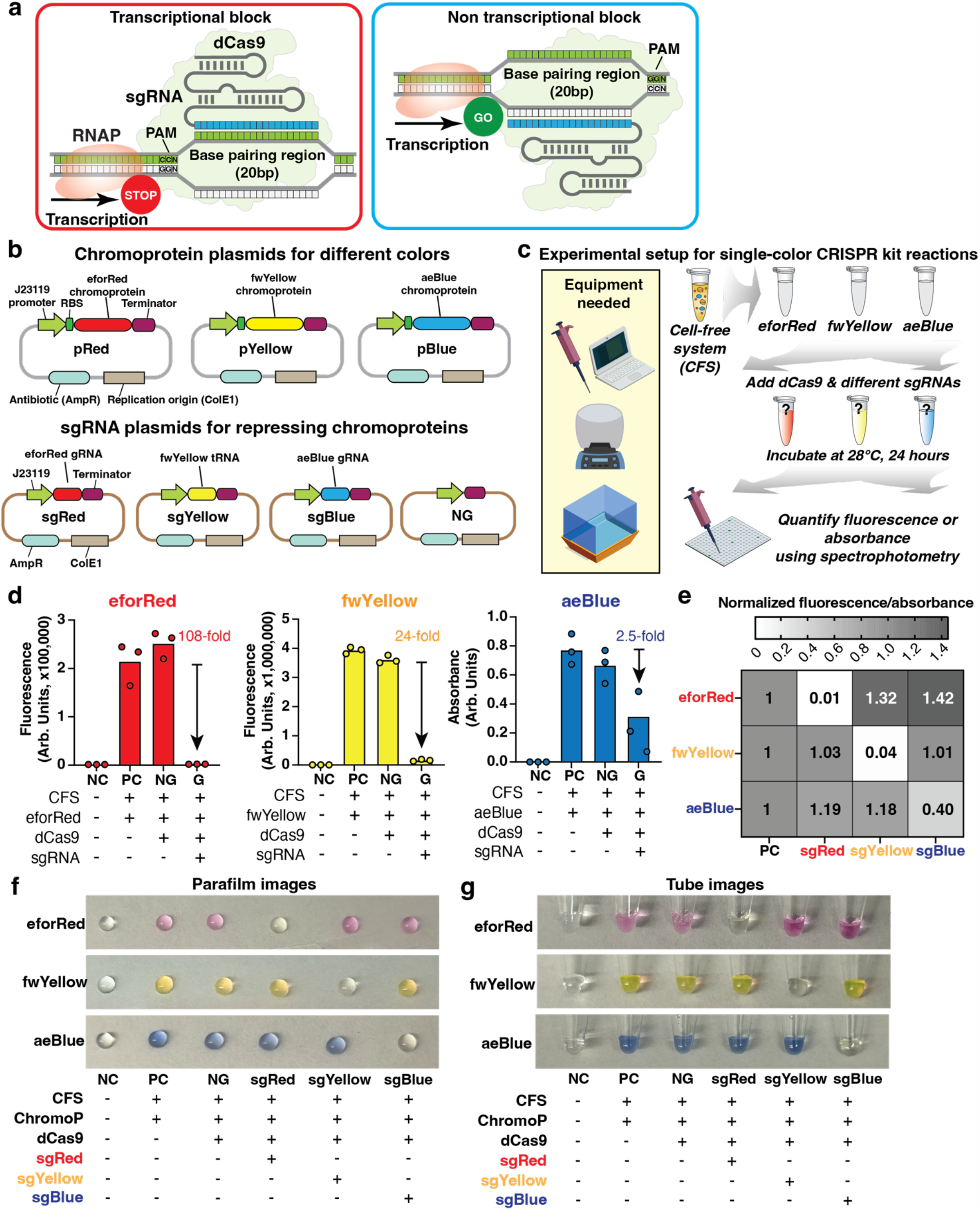
Design, implementation, and characterization of a single-color CRISPR kit under high-tech setup. **a**. Comparison of two designs of sgRNAs and how they alter transcriptional outcomes. On the left, the sgRNA is designed to target the non-template DNA strand (the DNA strand that is not used by RNA polymerase to make mRNA). In this case, the PAM is upstream of the sgRNA-DNA binding site and faces the incoming RNA polymerase. This configuration blocks transcription and leads to gene repression. On the right showing the sgRNA is designed to target the template DNA strand (the DNA strand that is used by RNA polymerase to make mRNA). In this case, the PAM is downstream of the sgRNA-DNA binding site and faces away from the incoming RNA polymerase. This configuration has been shown to be inefficient for gene repression. Our designed sgRNAs target the non-template DNA strand to maximize the gene repression effects. **b**. Schematic of plasmids that encode chromoproteins (top) and sgRNAs (bottom). For chromoproteins, we design three primary plasmids that encode eforRed, fwYellow, and aeBlue. Each plasmid contains a strong bacterial σ-70 promoter J23119, a strong ribosome binding site (RBS), codon-optimized coding sequence, and a transcriptional terminator. For sgRNAs, we design one sgRNA targeting each chromoprotein-encoding gene. Similarly, each plasmid contains a strong bacterial σ-70 promoter J23119, the sgRNA-coding sequence, and a transcriptional terminator. **c**. Experimental setup for the high-tech single-color experiment using the CRISPR kit. Briefly, the cell-free system is mixed with one chromoprotein plasmid in each tube and supplemented with one sgRNA plasmid. After incubation at 28°C for 24 hours, we quantify the fluorescence (for eforRed and fwYellow) or absorbance (for aeBlue) using a fluorescence plate reader. **d**. Experimental data showing the performance of high-tech single-color CRISPR kit reactions measured by a fluorescence plate reader. For each chromoprotein, we compare four conditions: negative control (NC) with water only, positive control (PC) containing CFS and the chromoprotein plasmid, no-guide condition (NG) containing CFS, the chromoprotein plasmid, dCas9 protein and an empty sgRNA plasmid (as shown in **Fig. 2b**), and targeting sgRNA condition (G) containing CFS, the chromoprotein plasmid, dCas9 protein and the cognate sgRNA. For each condition, three biological replicates with readings from the fluorescence plate reader are shown. The bars represent the mean. **e**. Heatmap showing the orthogonality of sgRNAs for repressing each chromoprotein expression. Normalized fluorescence (for eforRed and fwYellow) and absorbance (for aeBlue) to positive controls (PC) are shown. **f-g**. Phone images of the single-color CRISPR kit reactions transferred to a parafilm (**f**) or 0.6mL PCR tubes (**g**), after incubation at 28°C for 24 hours. From left to right, each group represent negative control (NC), positive control (PC, CFS + chromoprotein plasmid), no-guide (NG, CFS + chromoprotein plasmid + dCas9 protein + empty sgRNA plasmid), and targeting sgRNA for eforRed (sgRed), fwYellow (sgYellow), and aeBlue (sgBlue).

We tested whether different chromoproteins can be used in the cell-free CRISPRi experiments and whether they can be specifically repressed by their cognate sgRNAs. Using eforRed, fwYellow, and aeBlue chromoproteins encoded from plasmids as shown in **Fig. 2b**, we used pipettes to adddCas9 protein and sgRNA plasmids and incubated each reaction at 28°C for 24 hours in 1.5 mL Eppendorf tubes. We quantified fluorescence of eforRed and fwYellow or absorbance of aeBlue using a fluorescence plate reader after transferring samples to a 384-well plate (**Fig. 2c**, see **Methods**). We chose to measure absorbance of aeBlue because, unlike eforRed or fwYellow, there is not a suitable fluorescence excitation/emission wavelength that has been found for this protein^22^.

We observed strong chromoprotein expression without CRISPRi and efficient repression in the presence of dCas9 and cognate sgRNA for each chromoprotein (**Fig. 2d-e**). From the plate reader quantification, compared to the positive control (CFS with chromoprotein plasmids only), eforRed and fwYellow CRISPRi showed 108- and 24-fold of repression, respectively (**Fig. 2d**). On the contrary, aeBlue CRISPRi showed a smaller fold change (2.5-fold), likely due to the suboptimal measurement using absorbance^24^. Yet, visually from parafilm or tube images taken by smartphone, we observed strong pigmentation without CRISPRi and almost no color with CRISPRi (**Fig. 2f-g**). This suggests that phone images might offer a better method to quantify aeBlue expression level that is more consistent with the visual observations. We also confirmed that the sgRNAs acted orthogonally and only repressed their cognate targets (**Fig. 2e**). We noticed modestly elevated eforRed pigmentation in the presence of sgRNAs targeting feYellow or aeBlue, for reasons unknown (**Fig. 2e**).

In summary, our data suggests that the cell-free system provides an effective and robust way of performing gene-specific CRISPRi experiments with visible chromoprotein outcomes that can be quantified via the fluorescence/absorbance-measuring plate reader.

### Testing the performance of dual-color CRISPR kit in the laboratory setup

We next tested whether multiple chromoproteins can be used in the same reaction and whether each gene can be specifically repressed by their targeting sgRNAs. To do this, we used pipettes to mix eforRed and fwYellow plasmids with an equimolar ratio and added dCas9 protein and single or double sgRNA plasmids in 1.5 mL Eppendorf tubes (**Fig. 3a**). Similar to the single-color experiment, we quantified fluorescence of eforRed and fwYellow using the 384-well-based fluorescence plate reader (see **Methods**).

**Figure 3:**
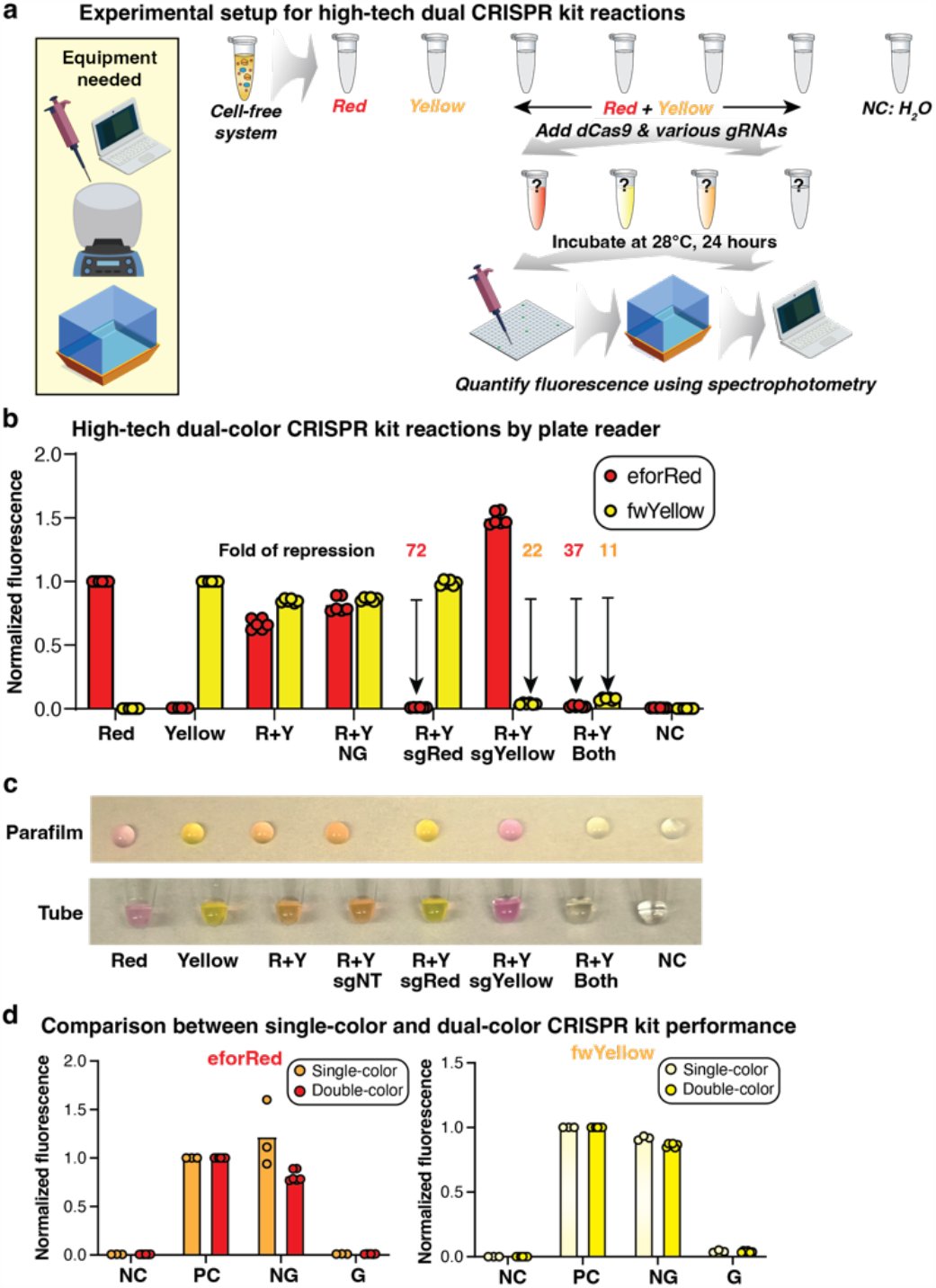
Design and characterization of dual-color CRISPR kit using the high-tech setup. **a**. Experiment setup for high-tech dual-color CRISPR kit reactions. The cell-free system is mixed with one or two chromoprotein plasmids with one or two sgRNA plasmids, incubated at 28°C for 24 hours, and the fluorescence is quantified using a fluorescence plate reader. **b**. Experimental data showing the performance of high-tech dual-color CRISPR kit reactions measured by a fluorescence plate reader. Each group of bars show eforRed or single-color positive controls, eforRed+fwYellow dual-color positive control, with an empty sgRNA plasmid (NG), eforRed-targeting sgRNA (sgRed), fwYellow-targeting sgRNA (sgYellow), and both eforRed- and fwYellow-targeting sgRNAs, and negative control (water only). Three biological replicates and six technical replicates (each biological replicates measured twice using the fluorescence plate reader) are shown. The bars represent the mean. **c**. Phone images of the high-tech dual-color CRISPR kit reactions on a parafilm (top) or PCR tubes (bottom), after incubation at 28°C for 24 hours. **d**. Comparison of chromoprotein expression between high-tech single-color and dual-color CRISPR kit reactions. The graphs show the normalized fluorescence for eforRed (left) and fwYellow (right) in single-color or dual-color reactions to their positive controls (CFS + chromoprotein plasmid). Three biological replicates are shown. For dual-color conditions, six technical replicates are shown. The bars represent the mean.

The dual-color CRISPRi experiment showed that individual chromoprotein genes can be specifically repressed by their individual sgRNA (**Fig. 3b**). When both sgRNAs were present, we observed strong repression of both chromoproteins. Both parafilm and tube images taken by smartphone showed visually distinct colors that were consistent with measurement by the fluorescence plate reader (**Fig. 3c**). Interestingly, the expression level of chromoprotein was highly comparable in the dual-color and single-color experiments (**Fig. 3d**). The level of CRISPRi-mediated repression was also similar when comparing the dual-color experiment with a single sgRNA to the single-color experiment: for eforRed, we observed 72-fold of repression in a dual-color experiment compared to 108-fold of repression in the single-color experiment; for fwYellow, it was 22-fold compared to 24-fold. The repression level was lower when using both sgRNAs in the dual-color expression (37-fold for eforRed and 11-fold for fwYellow). Nevertheless, this level of repression has generated distinct colors that can be easily discerned by the naked eye using phone images (**Fig. 3c**). Taken together, our experiment suggests that the dual-color CRISPR kit can be designed to generate visually distinct colors to offer facile readout.

### Designing and testing a frugal CRISPR kit by removing equipment requirements

Next, we designed a cost-effective and equipment-free CRISPR kit to increase its accessibility. We eliminated the need for specialized lab equipment by substituting lab equipment with accessible alternatives (**Fig. 4a**). We postulated that, instead of using pipettes and pipette tips, inoculation loops can be used to transfer liquid with volume around 1μL. To eliminate the incubator that provides accurate temperature and humidity control, we postulated that room temperature should be adequate to support sufficient cell-free protein production and CRISPRi activity. Furthermore, encouraged by our phone images taken for single-color and dual-color experiments, we hypothesized that a smartphone can be used in place of a fluorescence plate reader. Finally, to aid multi-color quantification using phone images, we developed an in-house algorithm and associated website for user-friendly analysis and data plotting.

**Figure 4:**
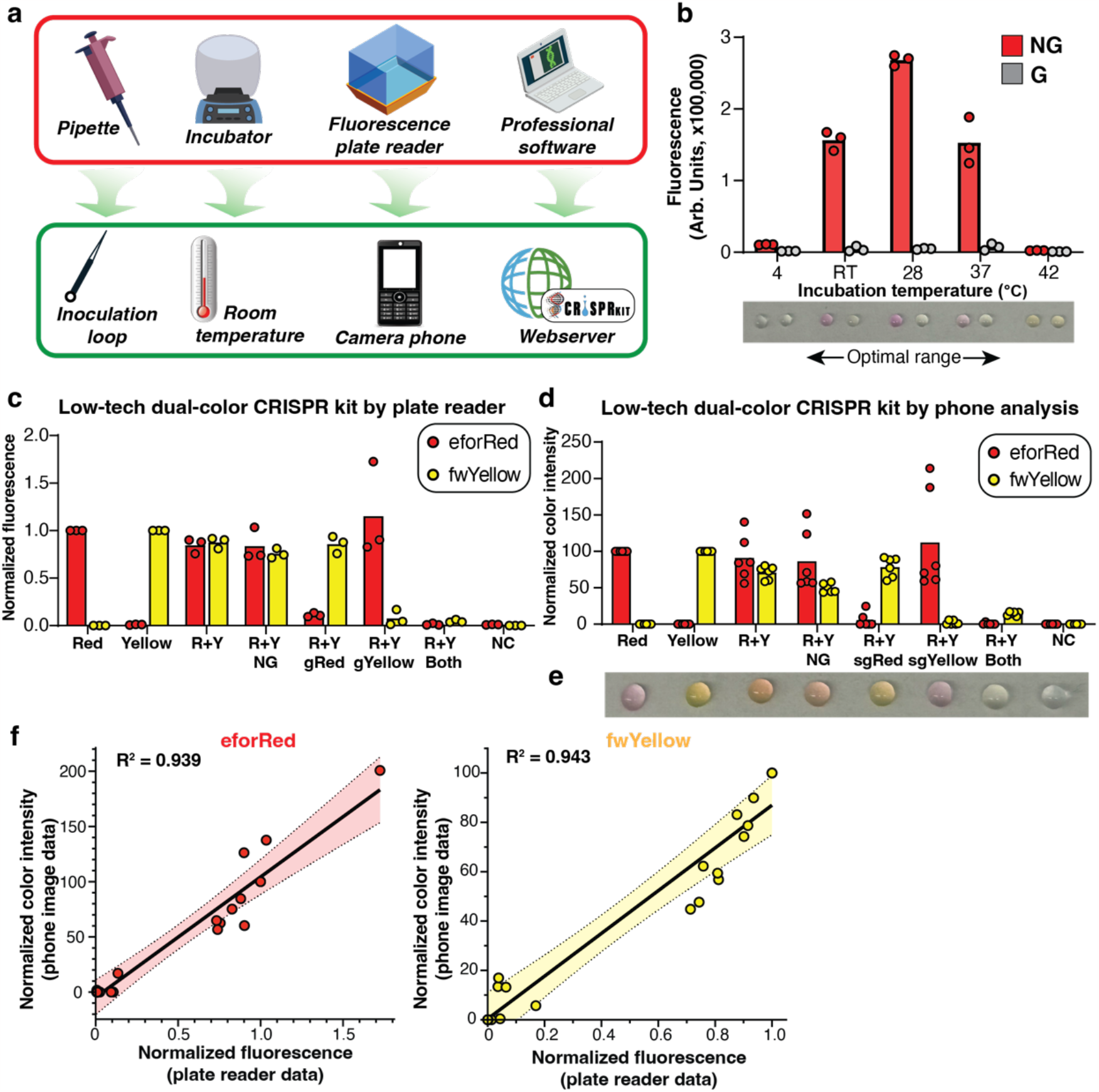
Design and characterization of frugal low-tech dual-color CRISPR kits. **a**. Schematic showing replacing laboratory apparatus and equipment with frugal alternatives. Specifically, we replace pipettes with inoculation loops, incubator with a room temperature setup, fluorescence plate reader with a smartphone, and professional analytical software with our web server for phone image data analysis. **b**. Characterization of the performance of the single-color CRISPR kit at different temperatures using eforRed. Five temperatures are tested, ranging from 4°C to 42°C, representative of ambient temperatures. All reactions are incubated for 24 hours. Three biological replicates are shown for each group and the bars show the mean fluorescence. Red, non-targeting sgRNA; gray, targeting sgRNA. On the bottom, a representative phone image of CRISPR kit reactions transferred to a parafilm is shown. **c**. Experimental data showing low-tech dual-color CRISPR kit reactions measured by a fluorescence plate reader. From left to right, each group of bars (red – eforRed, yellow – fwYellow) show eforRed single-color positive control, fwYellow single-color positive control, eforRed + fwYellow dual-color positive control, dual-color with empty sgRNA plasmid (NG), dual-color with eforRed-targeting sgRNA (sgRed), dual-color with fwYellow-targeting sgRNA (sgYellow), dual-color with both eforRed- and fwYellow-targeting sgRNAs, and negative control (water only). For each condition, three biological replicates are shown. The bars represent the mean. **d**. Experimental data showing low-tech dual-color CRISPR kit reactions measured by a smartphone and analyzed by our algorithm. The order of the reactions follows the same as **c** for side-by-side comparison. For each condition, three biological replicates are shown. The bars represent the mean. **e**. A representative phone image of the low-tech dual-color CRISPR kit reactions transferred to a parafilm, after incubation at room temperature for 24 hours. The order of the reactions follows the same as shown in **c. f**. 2D scatter plots showing comparison of fluorescence plate reader data (x-axis) and phone image data (y-axis) for eforRed (left) and fwYellow (right). Individual dots represent biological replicates in each condition as shown in **c** and **d**. Black lines represent the linear fit and the dotted lines represent 99.9% confidence interval. R^2^ values are shown in the diagrams for the linear regression.

We first performed an experiment to verify whether room temperature (RT, ∼22 °C) enabled efficient chromoprotein expression and CRISPRi activity using the cell-free system (**Fig. 4b**). Using eforRed, we quantified the chromoprotein expression (without sgRNA) and repression (with sgRNA) using the fluorescence plate reader. Our data suggests the optimal temperature range to carry out the CRISPRi cell-free reaction is around 28 °C. Notably, room temperature and 37 °C also showed strong expression and repression. On the contrary, 4 °C and 42 °C greatly diminished both protein production and CRISPRi activity. Therefore, we confirmed room temperature offers an acceptable condition for the CRISPR kit, thus eliminating the need for an incubator.

We next developed an algorithm, termed CRISPectra, to quantify color intensity, as a proxy for chromoprotein expression level, using the high-tech single-color and dual-color experimental data (see **Methods**). The CRISPectra algorithm extracts the RGB (red-green-blue) values from individual pixels, averages the values of all pixels in a selected area in the image, and outputs normalized color intensity values for each reaction by comparing with the average RGB values of single-color reactions. Our results showed that the algorithm and phone images generated data for eforRed and fwYellow consistent with those measured by a fluorescence plate reader (**Fig. S1a**). Interestingly, our phone image method showed better repression for aeBlue, which was more aligned with the visual observation.

Comparing the phone image data with plate reader data, we observed high correlation between samples for all three chromoproteins (**Fig. S1b**). We further characterized dual-color high-tech experiments using the algorithm and phone images and observed similarly highly correlation between phone images and plate reader measurement (**Fig. S2a**). In summary, our data suggests that phone images and the CRISPectra algorithm provide a reasonable approximation of chromoprotein expression as an alternative to the fluorescence or absorbance measurement by the plate reader.

### Characterization of the frugal CRISPR kit performance in a laboratory setting

We next performed a dual-color CRISPRi experiment in a frugal setting, using inoculation loops to transfer liquids and incubated the reactions at room temperature for 24 hours (**Fig. 4a**). Using eforRed and fwYellow chromoproteins, our data demonstrated strong and consistent CRISPRi repression of cognate chromoproteins in the presence of individual or double sgRNAs, measured by the fluorescence plate reader (**Fig. 4c**) or phone images (**Fig. 4d**). The reactions using the frugal setup showed easily visible and distinct colors (**Fig. 4e**). We also compared plate reader data with phone imagery data quantified using the CRISPectra algorithm. We saw strong correlation between the two methods (R^2^ = 0.939 and 0.943 for eforRed and fwYellow, respectively, **Fig. 4f**). This experiment confirmed that the frugal CRISPR kit design and the quantification method reliably reported CRISPRi activity without the need for equipment.

### Real-world testing of the frugal CRISPR kits with high school students

We next tested whether the frugal CRISPR kit can be used by high school students in a non-laboratory frugal setting (**Fig. 5a**). We manufactured the frugal CRISPR kits in bulk and distributed them to high school students (**Fig. S3a-b**). Data from nine experiments performed by high school students showed that eight experiments generated consistent chromoprotein expression and CRISPRi activity (**Fig. 5b**), despite one experiment failing (data not shown). Nevertheless, the colors from the samples in most experiments were easily visible and discernible. Using the CRISPectra algorithm, we quantified the performance of eight experiments and compared the phone image results with those measured by the plate reader for each experiment (**Fig. 5c** and **Fig. S4**). Our data showed that the frugal setup generated results consistent with those measured by the plate reader, although with a higher variation that was partly due to the lighting condition used by the students to take photos (**Fig. 5c**). The performance of these experiments was also comparable but more variable than those performed in a laboratory setup (**Fig. 4d**). We attributed the variation to the accuracy of using inoculation loops in liquid transfer by the students.

**Figure 5:**
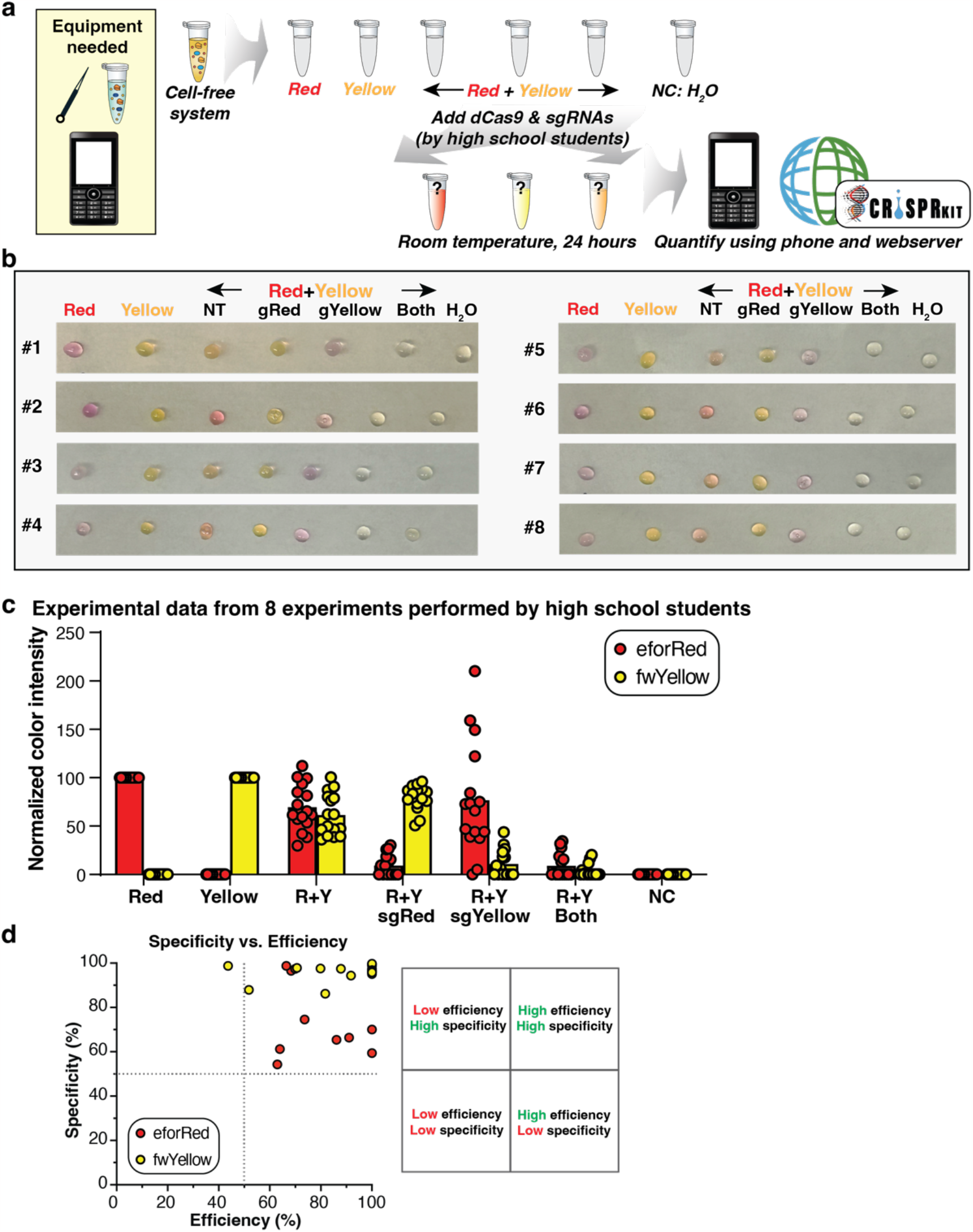
Testing the frugal dual-color CRISPR kit by high school students in classrooms. **a**. Experimental setup of the frugal dual-color CRISPR kit in a non-equipment setting. The reagents are provided to the students, together with a detailed stepwise protocol. The students transfer the reagents including chromoprotein plasmids, dCas9, and sgRNA plasmids into reaction tubes preloaded with the cell-free system. The reactions are left at room temperature for 24 hours. Smartphones are used to image the reactions and uploaded to a web server for image analysis. **b**. Phone images of frugal dual-color CRISPR kit reactions (#1 to #8) on a parafilm. Eight sets of data were shown. From left to right, each group of bars show eforRed or fwYellow single-color positive control, eforRed+fwYellow dual-color positive control, with the eforRed-targeting sgRNA (sgRed), with the fwYellow-targeting sgRNA (sgYellow), with both sgRNAs, and negative control (water only). For each condition, eight experimental replicates (each data point measured twice as technical replicates of phone image analysis) are shown. Bars represent the mean. **c**. Experimental data showing frugal dual-color CRISPR kit reactions measured by smartphones. **d**. 2D scatter plot showing the calculated efficiency and specificity for each reaction for eforRed (red) and fwYellow (yellow). The top right quadrant shows reactions with high efficiency and high specificity.

We reasoned that teaching students the concept of efficiency and specificity helps to better understand the core concepts of CRISPR experiments. Therefore, it is important to report the efficiency (the level of repression on the target gene) and specificity (the level of crosstalk repression on another gene) for CRISPR experiments. We calculated the efficiency and specificity of each experiment (**Fig. S5**). Our results showed high efficiency and specificity among experiments performed by high school students (**Fig. 5d**).

### Expanding the frugal CRISPR kit as a versatile and modular educational module for genetic engineering in the high school setup

We next expanded the utility of CRISPR kits towards hypothesis-driven experiments in the classroom. We reasoned that combinations of different chromoproteins provide numerous outcomes that can be directly observed without equipment, therefore inspiring the students to formulate and execute their own plans using a streamlined frugal CRISPR kit.

We first streamlined the design of our kits to minimize its components (**Fig. 6a**). We also provided detailed manual, protocol, and learning contents (**Supplementary Information**). We next tested combinations of two or three chromoproteins using the cell-free system but without CRISPR components. Our results showed that chromoprotein combinations can generate a diverse panel of visible colors, forming a basis for using the CRISPR kit to knock down single or multiple chromoprotein genes and measuring different outcomes (**Fig. 6b**). This offers an opportunity for students to make their own designs, in a ‘guess-and-test’ manner, to figure out the composition of genes in a tube that is provided to them by the teacher (**Fig. 6c**).

**Figure 6:**
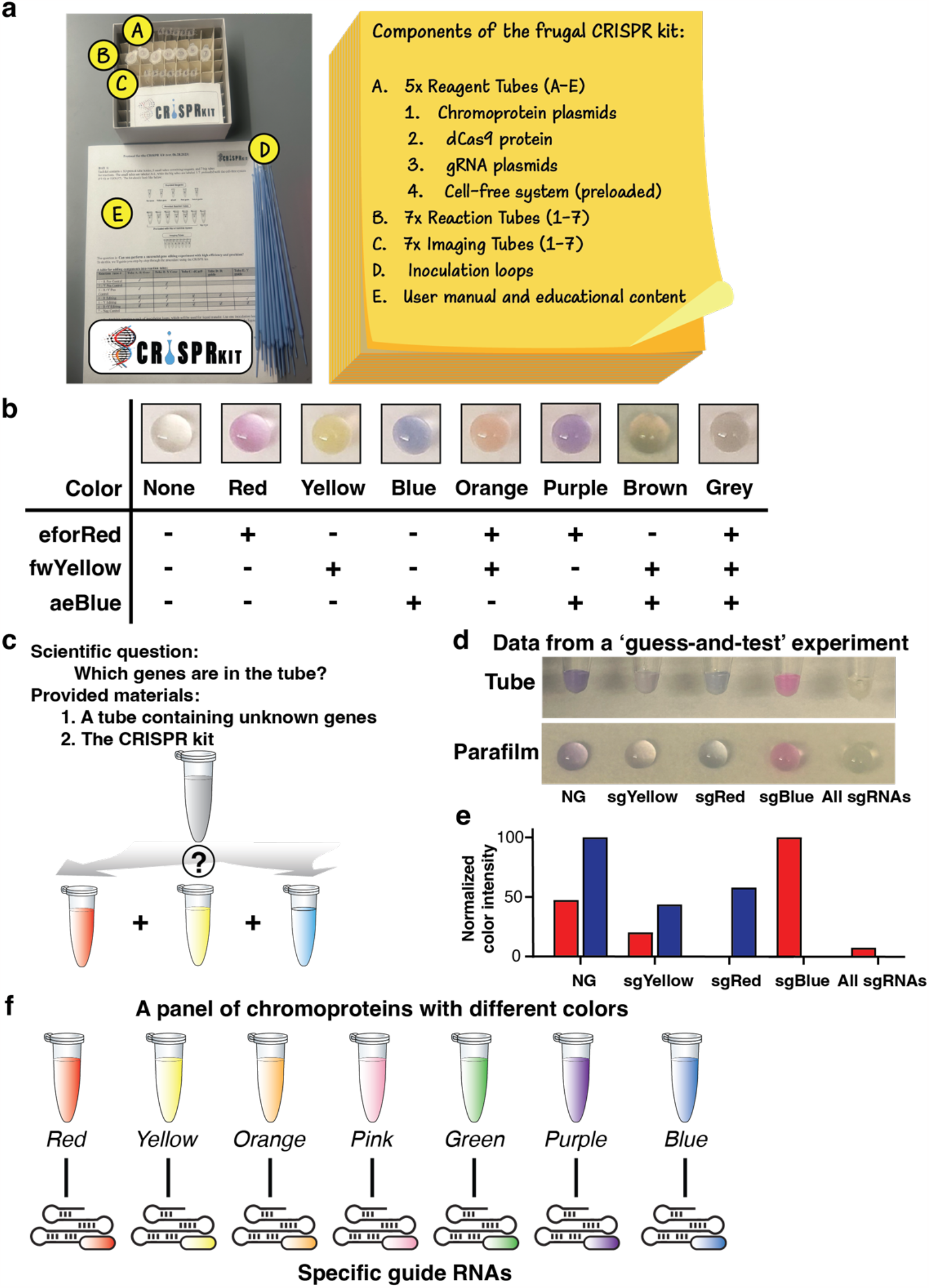
Expanding the frugal CRISPR kit into a modular laboratory module for open-endedness teaching. **a**. Illustration of the components of the frugal CRISPR kit with a phone image (left) and a list (right). A, Reagent tubes (5), containing the chromoprotein plasmid for eforRed, fwYellow, dCas9 protein, and sgRNA targeting eforRed or fwYellow. B, Reaction tubes (7, preloaded with the cell-free system). C, Imaging tubes (7), for the phone image purpose. D, Inoculation loops (17 loops for each kit, and 3 spare ones). E, User manual and protocol. **b**. Phone images of combinations of different chromoproteins in the cell-free system reactions transferred to a parafilm. **c**. Schematic of a ‘guess-and-test’ experiment. The question asked is which genes are in the tube? A tube containing unknown plasmids encoding one or more chromoproteins and the frugal CRISPR kit are provided to the students. **d**. Experimental data showing results using the CRISPR kit reactions to demystify the genes in a tube. **e**. Quantification of experimental data using CRISPectra algorithm. **f**. Schematic showing expanding the frugal CRISPR kit to a variety of chromoproteins with different colors and specifically targeting sgRNAs for more possibilities of ‘guess-and-test’ experiments.

We designed an experiment, by providing a mixture of eforRed and aeBlue plasmids to students. In this experiment, the students were unaware of which genes were in the tube but instructed to test their own hypothesis using the CRISPR kit that contained the cell-free system and each of the sgRNAs that target eforRed, fwYellow, and aeBlue (**Fig. S6**). The student made an experimental design by adding the mixture of plasmids to the cell-free system and different conditions of sgRNAs that target either fwYellow, eforRed, aeBlue, or all of them. After incubating at room temperature for 24 hours, the student took photo images as shown in **Fig. 6d**, quantitatively analyzed the results using the CRISPectra algorithm (**Fig. 6e**) and reported the unknown genes in the tube were a mixture of red and blue genes.

We reasoned that there exists a panel of diverse chromoproteins that can be harnessed to build an expanded CRISPR kit, each with a specific targeting sgRNA, to enable more possibilities of such ‘guess-and-test’ experiments (**Fig. 6f**). We believe such experiments offer a cost-effective way for high school students to perform their own designed CRISPR experiments, thus promoting their learning and inspiring their interests towards biotechnology. Finally, we designed a user-friendly website (https://crisprkit.org) containing information of the CRISPR kit, protocols, and analysis algorithm, to help disseminate the kits among broad communities (**Fig. S7**).

## Discussion

In this study, we designed and implemented a safe, frugal, cost-effective CRISPR kit that can be broadly distributed to high school students to gain hands-on experiences on CRISPR experiments. Our CRISPR kit eliminates the requirements for equipment and can be essentially performed in any environmental setting (e.g., classroom or garage). We chose to use CRISPRi with a nuclease-dead dCas9, instead of the cutting enzyme Cas9, to minimize potential impact on the environment if these reagents were disposed of because dCas9 does not induce permanent DNA alterations. Our results in the laboratory setting showed that the frugal kits without using pipettes or equipment performed comparably well as the CRISPR reactions carried out using standard laboratory equipment. We also developed an easy-to-use computational algorithm, CRISPectra, to aid the quantification of the chromoprotein expression based on pigment color intensity. Our real-world tests confirmed the reliability of the CRISPR kit when used by high school students who had no prior experience in CRISPR. These tests together suggest that the CRISPR kit offers an easy and reliable solution to aid hands-on experiment experiences to high school students.

Democratizing biotechnology has the potentially to extend the benefits of bioengineering advancements to a diverse population. Nonetheless, this goal has faced challenges due to a scarcity of innovations and useful tools. Equitable and accessible education is crucial to promote biotechnology democratization, which urgently needs the development of tools that are safe, accessible, and cost effective. CRISPR technology is easy to understand, has broad impact on the real world, and has gained tremendous exposure in the past decade. We envision that the frugal CRISPR kit can be manufactured in large quantities and distributed to high school teachers, who can use them to prepare for their teaching. A freezer was all needed for the short-term storage of reagents at 4°C and long-term storage at -20°C. We anticipate that the components can be freeze dried to enable room-temperature storage in the future. From our interactions with high school teachers and students, despite a strong interest in learning about the CRISPR technology, there is a lack of effective hands-on education tools to perform safe and affordable CRISPR experiments in high school classroom settings. We believe the frugal CRISPR kit developed here can eliminate major financial barriers and allow teachers and students to easily adopt for their teaching and learning purposes.

Compared to other reported cell-free systems using CRISPR^11,13,14,26^, our kit eliminates the need for equipment (e.g., pipettes, incubator, and imager), relies solely on smartphones for measurement, and an in-house CRISPectra algorithm for quantification, which increases accessibility to students. Our current CRISPR kit makes use of the commercial transcription-translation system and all reagents needed to perform the frugal CRISPR kit experiments can be ordered easily from retailers. We foresee that, with advances of the cell-free system with higher robustness and affordability, the cost of the CRISPR kit can decrease dramatically (presumably lower than $1 per kit).

From our analysis of successful and failed frugal CRISPR kit experiments, the performance of the kit is mainly dependent on the quality of purified plasmid DNA and the accuracy of liquid transfer using inoculation loops. For example, we noticed drastically different performance of chromoprotein production using suboptimally purified chromoprotein plasmids. The failed real-world experiment using CRISPR kit most arise from the failure to transfer liquid from the reagent tubes to reaction tubes. For phone images, considering that the lighting and shading in the photos may play a role affecting the data quantification, we’ve designed the algorithm to minimize the lighting impacts. In the future, we plan to incorporate diverse chromoproteins with more color options and characterize their specific sgRNAs. We also envision the incorporation of CRISPR activation (CRISPRa) in parallel to orthogonal CRISPRi for genetic circuitry-type CRISPR kits to expand the system into a synthetic biology system. We believe that affordable access to revolutionary CRISPR technology is a valuable tool, with which educators and students can pursue their interests in life sciences and bioengineering, thus contributing to equitable and accessible education regardless of their socioeconomic background.

## Methods

### Plasmid cloning and preparation

All primers were designed in Benchling and synthesized by Stanford’s Protein and Nucleic Acid Facility. Gibson assembly was used for the construction of pSLQ3533 (eforRed plasmid) and pSLQ3534 (fwYellow plasmid). To clone the chromoprotein plasmids, pSLQ220 was digested by NheI and XhoI. gBlocks encoding the fwYellow and eforRed genes were ordered from IDT. gBlocks and the digested vector were joined using an NEBuilder HiFi DNA Assembly kit during a 1 hr incubation at 50 °C. Plasmid DNA from the Gibson assembly reaction was then transformed into Stellar Competent Cells (Takara Bio).

In-Fusion cloning (Takara Bio) was used for the construction of plasmid pSLQ14101 (aeBlue plasmid). pSLQ3533 was digested by restriction enzymes BglII and XhoI. Plasmid encoding the aeBlue gene was ordered from Addgene (#117846). The aeBlue chromoprotein gene was amplified in a PCR reaction using a KAPA Hi-Fi kit. Following electrophoresis, PCR products were extracted from a gel using a NucleoSpin Gel and PCR Cleanup kit (Machery-Nagel). Purified PCR products were inserted into the digested backbone using an In-Fusion reaction. Plasmid DNA from the In-Fusion reaction was then transformed into Stellar Competent Cells.

To design sgRNAs, we selected a 20-bp sequence with a nearby PAM sequence (NGG) on the coding region of each chromoprotein as the target site and used algorithm CRISPR-ERA (http://CRISPR-ERA.stanford.edu) to verify their specificity. For the cloning of pSLQ3538 (eforRed-targeting sgRNA plasmid), pSLQ14103 (aeBlue-targeting sgRNA plasmid), and pSLQ14107 (fwYellow-targeting sgRNA plasmid), forward and reverse primers for a full-plasmid PCR were designed. Forward and reverse primers contained 15 base pair regions homologous to each other for joining in an In-Fusion reaction. The forward primers contained a region encoding the desired spacer sequence for the crRNA. pSLQ3529 was used as the DNA template for a full-plasmid PCR using the described primers, which was performed using a KAPA Hi-Fi kit. Following electrophoresis, PCR products were extracted from a gel using a NucleoSpin Gel and PCR Cleanup kit. Purified PCR products were circularized using an In-Fusion reaction. Plasmid DNA from the In-Fusion reaction was then transformed into Stellar Competent Cells.

All constructs were selected on LB-agar supplemented with 100 ug mL^-1^ carbenicillin. Sequence-verified clones were purified using a Plasmid Plus Midi Kit (Qiagen), then purified again using a Nucleospin Gel and PCR Clean-Up Mini kit (Machery-Nagel). The final purification step is essential to ensure optimal performance of plasmids in myTXTL Sigma 70 Master Mix.

### High-tech CRISPR kit reaction

All fluid transfer was performed using micropipettes. MyTXTL Sigma 70 Master Mix was purchased from Daicel Arbor Biosciences and vortexed and spun down in a centrifuge before each experiment. 9 uL of myTXTL Sigma 70 Master Mix was allotted into 1.5 mL Eppendorf tubes. All other reaction components, if included in a reaction, were added to the following target concentrations: 2.1 ng uL^-1^ dCas9 V3 protein (IDT, cat no. 1081066); 5 nM chromoprotein plasmid DNA; 2.5 nM sgRNA plasmid DNA. Water was added to bring the final reaction volume to 15 uL. Reactions were incubated at 28°C for 24 hours.

### High-tech chromoprotein quantification

5 uL of each sample was pipetted into a Hard-Shell 384-well clear microplate (Bio-Rad #HSP3805), and fluorescence and absorbance readings were taken by a Synergy H1 microplate reader (BioTek). The excitation and emission wavelengths used to quantify fluorescence of eforRed and fwYellow were as follows: eforRed ex 589 nm, em 615 nm; fwYellow ex 520 nm, em 550 nm. The absorbance wavelength used to quantify the absorbance of aeBlue was 593 nm.

### Low-tech CRISPR kit reaction

10 uL of myTXTL Sigma 70 Master Mix was allotted into 1.5 mL Eppendorf tubes. All other reaction components, if included in a given reaction, were added to the following target concentrations: 2.1 ng/uL dCas9 V3 protein (IDT); 5 nM chromoprotein plasmid DNA; 2.5 nM sgRNA plasmid DNA. Reagent stocks were diluted such that adding 1 uL of each to a given reaction would achieve the target concentration. For the addition of each reaction component, a 1 uL inoculation loop was first inserted into the stock solution and twirled several times while fully submerged. Then the loop was removed and checked by eye to ensure fluid was present within the loop. The loop was then inserted into the reaction mixture and twirled several times while fully submerged. Loops were then rinsed with running water and dried with a paper towel for later reuse. Enough water to bring the final reaction volume to 15 uL was pipetted into each reaction. Reactions were incubated at room temperature for 24 hours.

### Image acquisition and pre-processing for smartphone-based data analysis

#### Sample Preparation

For image acquisition, 4 µL aliquots from each reaction tube were sequentially transferred onto parafilm.

#### Image Acquisition

All experimental images were captured using an iPhone 14. Each image was taken in the same laboratory space equipped with consistent and optimal lighting conditions. The imaging angle was maintained at a slightly slanted angle relative to the parafilm’s surface at approximately an arm’s length. This enabled the concealment of shadows to mitigate glare.

#### Image Segmentation and Pre-Processing

After image acquisition, the images were manually cropped in a rectangular shape within the circular regions of the liquid droplets on the parafilm. Once segmented, the images were either retained in their original state or subjected to a minimum filter using the FIJI (ImageJ) software platform if image glare is deemed too significant. The minimum filter operates through grayscale erosion, replacing each pixel in an image with the smallest pixel value in its proximate neighborhood. This step is particularly vital in instances where an image manifests pronounced glare, as the filter effectively attenuates prominent bright spots. For images necessitating this filter, a radius equivalent to half of the image’s pixel width was selected.

#### Expression Analysis

Following the pre-processing stage, all the images were then fed into the phone-data analysis algorithm CRISPectra for color intensity expression analysis.

### CRISPectra Algorithm – Phone-Based Data Analysis Pipeline for CRISPR

#### 1. RGB-Value Extraction

Each image represents a distinct experimental condition. The primary RGB (Red, Green, Blue) values were extracted from these images, and their mean values *u*_*r*_, *u*_*g*_, *u*_*b*_ computed using:

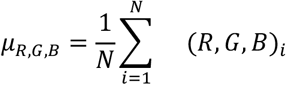

where N is the total number of pixels in the image.

#### 2. Negative Control Normalization

There were two foundational controls in each experiment: the Negative Control (NC) and the Positive Control (PC). The NC was established using only water, serving as a baseline, while the PC incorporated solely the chromoprotein gene, functioning as a benchmark for maximum intensity. Beyond these controls, the experimental conditions included targeting sgRNA (G) and no sgRNA (NG), capturing the varying dynamics of CRISPR interactions.

To account for non-specific signals and background noise, RGB values across all conditions were normalized against those obtained from the Negative Control (NC) condition. Specifically, for each condition *i* and its respective channel *j* (R, G, B), the values were adjusted using:

Let:

- *i* be the index for the condition (e.g., NC, PC, condition 3, 4 …)
- *j* represent the channel (R, G, B)

With this, the adjusted value for a specific channel in a specific condition can be given as:

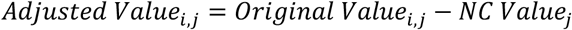

Where:

- *Original Value*_*i,j*_ is the value for the *j*^*th*^channel (R, G, B) for the *i*^*th*^ condition.
- *NC Value*_*j*_ is the value for the *j*^*th*^ channel in the NC condition.

#### 3. Dimensional Reduction

Next, we employed dimensional reduction on the RGB values of each experimental condition. This transformed the 3-dimensional RGB data of each condition into a singular representative value, thus condensing the color information into a quantifiable metric.

Our paper primarily presents two types of experiments. The first involves a singular chromoprotein, focusing on measuring the exclusive expression of this protein. The second delves into dual color experiments, which included reactions with two distinct chromoproteins. This necessitated the extraction of intensity levels for both colors. We will now explain how the computation applied to both types of experiments.

##### For Single Color

When the experiment involved only one chromoprotein of color A, the reference vector, *Vector*_*A*_, was derived from the RGB values of the PC condition, which corresponded to the condition where only color A was present in the cell-free mix. Mathematically, the magnitude of this vector is given by:

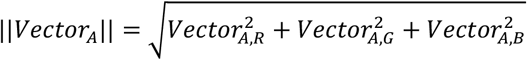

Where:

- 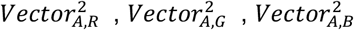 represents the components of the PC *Vector*_*A*_ in the red, green, and blue channels, respectively.

For instance, within the context of the paper’s single-color experiments, consider an experiment with four conditions: NC, PC, NG, and G. After the initial step of normalizing each of the RGB values with the NC values, each condition still retained three RGB values. The dimensional reduction step was thus implemented to transform the three-dimensional data into one singular, representative value for each condition. The mathematical representation of the projection is as follows:

For each condition *i*:

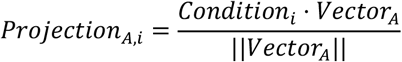

Where:

- *Projection*_*A,i*_ represents the singular value describing the intensity level of color A in condition *i*.
- Condition *i* represents the RGB values of the *i*^*th*^ condition, which are then subjected to a dot product operation with the RGB values of the positive control *Vector*_*A*_.

Since there were 4 conditions, 4 projections were made to transform the dataset from its original 12 values—attributed to 3 RGB components across 4 conditions—to just 4 distinct values. Each of these values corresponded uniquely to one of the conditions: NC, PC, NG, and G, providing a representation of the intensity level of the color A chromoprotein in each respective condition.

##### For Dual Color

When the experiment involved two chromoproteins of color A and color B, we independently extracted the intensity levels for each chromoprotein. A two-step projection methodology is implemented, where each color was computed separately.

Same as the single-color experiments, for the first chromoprotein color A, the reference vector, *Vector*_*A*_, was derived from the RGB values of the PC condition for color A.

We also computed the second reference vector, *Vector*_*B*_, derived from the RGB values of the PC condition for the second chromoprotein - color B. The two projections are mathematically defined as:

For each condition *i*:

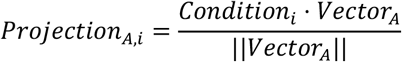

And

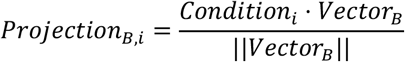

This allowed for the extraction of two unique intensity values for each condition from its original RGB values, one for each of the two chromoproteins. For instance, within the context of the paper’s dual-color experiments, consider an experiment with 7 conditions: NC, PC (color A only), PC (color B only), PC (color A + color B), *G*_*A*_ (targeting color A),*G* (targeting color B), and *G*_*A*+*B*_ (targeting both color A + color B). The projections generated a total of 14 unique values: a set of 7 for each condition for color A, and 7 for each condition for color B.

### 4. Crosstalk Minimization

In dual-color experiments, color overlap between different chromoproteins could lead to inaccurate intensity readings. To correct this, we calculated the shifts for each color and adjusted their projected values accordingly.

In the context of single-color experiments, where only one chromoprotein is present, this adjustment step was not necessary.

#### Calculation of Shifts

Color A shift per Color B value:

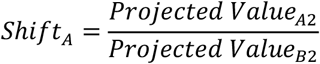

*Projected Value*_*A*2_ denotes the projection value of PC (color B only) for color A, while

*Projected Value*_*B*2_ denotes the projection value of PC (color B only) of color B.

Color B shift per Color A value:

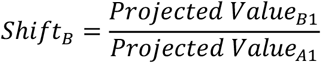

*Projected Value*_*A*1_ denotes the projection value of PC (color A only) for color A, while

*Projected Value*_*B*2_ denotes the projection value of PC (color A only) of color B.

## Adjustment of Projected Values

Using these shift values, we then calculated adjusted values for color A and color B, mathematically defined as:

For each color A condition *i*:

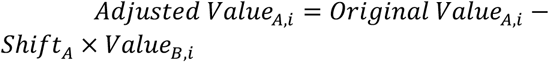

For each color B condition *i*:

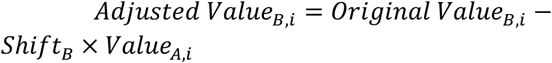

These adjustments subtracted the influence of one color from the other, ensuring that the resulting values purely represented each chromoprotein’s intensity level. This calibration ensures that when assessing the intensity of a specific chromoprotein, the projection value attributed to the contrasting color was nullified. For instance, when evaluating Color A’s projection intensity values for the Color B PC (color B only) condition, its entry is 0, and vice versa.

In the final step, for both single and dual color experiments, all derived values were normalized to the respective color’s PC value. This normalization effectively expressed each condition as a percentage relative to the PC’s intensity level. As a result, the PC assumed a value of 1, with all other condition values being presented as ratios in relation to this benchmark.

### 5. Graphical Representation and Adjustments

Upon obtaining the normalized values, additional adjustments were made to enhance the clarity and interpretability of the plots:

#### Scaling for Clarity

All values were multiplied by 100. This scaling ensured that the NC condition had a value of 0, with the PC peaking at 100, offering a more intuitive percentage-based interpretation of the results.

#### Negative Value Treatment

To better reflect biological reality, any values that became negative after adjustments were reset to 0. This adjustment ensured that the graphical depiction was consistent with the understanding that a negative value is biologically implausible for gene expression.

#### Plotting

The graphs were then plotted on Prism 9 (GraphPad) using these adjusted values for all phone imagery data.

#### Correlation of Phone-Based and Plate Reader Data

To assess the efficacy of the phone-based method of measuring color intensity in approximating gene expression levels, its results were benchmarked against data obtained from a high-tech plate reader through correlation analysis.

#### Data Plotting

The collected data sets were plotted on a scatter plot using DataGraph (Visual Data Tools). Each point on this plot represents an experimental condition, with its X-coordinate denoting the plate reader value and its Y-coordinate indicating the phone-based value.

#### Regression and Correlation Analysis

A linear regression analysis was performed on the plotted data to generate the line of best fit. The strength and direction of the linear relationship between the two datasets were quantified using the *R*^2^ (Pearson correlation coefficient) value. A value close to 1 indicates a strong positive correlation between the phone-based and plate reader data, suggesting the phone-based method’s robustness. The confidence interval is also shown, providing a range within which the true regression line is likely to fall, indicating the precision of the regression estimate.

#### CRISPRi Gene Repression Efficiency and Specificity Analysis

To assess the performance of the CRISPRi system in terms of its gene repression capability for dual color experiments, two key metrics were plotted: Efficiency and Specificity.

##### Efficiency

The efficiency metric reflects the ability of the CRISPRi system to selectively target and repress gene expression. It is calculated based on the reduction in the expression of the targeted gene compared to a condition where both genes (Red and Yellow) are expressed.

**For Yellow:**

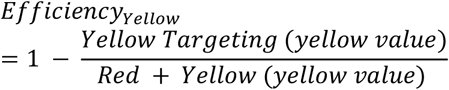

**For Red:**

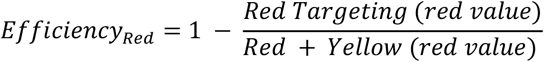

##### Specificity

Specificity provides insight into the system’s ability to selectively target one gene without affecting the other. A high specificity indicates minimal off-target effects.

**For Yellow:**

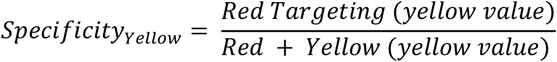

**For Red:**

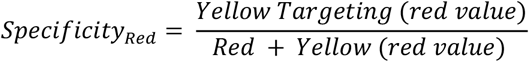

## Author Contributions

M.L. and L.S.Q. conceived of the original idea. M.C.C., M.L., M.L.R. and L.S.Q. contributed significantly to the frugal design. M.C.C., M.L., and M.L.R. cloned the chromoprotein plasmids and sgRNA plasmids. M.C.C. and M.L. performed the high-tech experiments. M.C.C. performed the low-tech experiments. M.L. developed the computational algorithm and the website. M.C.C. analyzed the experiments performed using laboratory equipment. M.L. analyzed experiments performed using smartphone images. W.M. and A.S. performed real-world tests. W.M. made significant improvements to the design and imaging conditions. M.C.C., M.L. and L.S.Q. plotted the figures. L.S.Q. secured funding. L.S.Q. wrote the manuscript with input from all authors.

## Acknowledgements

The authors thank Emily Ma for testing the CRISPR kit and providing valuable advice on improving the kit design. The authors thank Prof. Huijun Zhou Ring and Daniel Stauber for helpful discussions. The authors thank Dr. Ross Venook, Dr. Alex Engel, and Dr. Phillip Kyriakakis for organizing undergraduate students at Stanford University for testing the CRISPR kit and offering helpful advice. The authors thank high school educators and teachers, including Meghan Strazicich, James Stiltner, Temy Taylor, I-Heng McComb, and Johnson Huynh, for testing the kits and offering valuable advice. The authors thank Prof. Michael Jewett and Dr. Peivand Sadat Mousavi for facilitating cell-free experiments. The authors thank Sa Cai for testing the kit. The authors thank all members in the Lei Stanley Qi lab for helpful discussions. L.S.Q. acknowledges support from the National Science Foundation CAREER award (no. 2046650). L.S.Q. is a Chan Zuckerberg Biohub – San Francisco Investigator. The work is supported by NSF CAREER award and partly by the making@Stanford initiative (making.stanford.edu).

## Data and Code Availability

All plasmids encoding the CRISPR kit will be available on Addgene: https://www.addgene.org/Stanley_Qi/. The CRISPR kit can be requested from the CRISPR kit website at https://www.crisprkit.org. The code for data analysis is available via Github at https://github.com/matthewbhlau/crispr-kit/tree/main.

## Supplemental Information for

## Supplemental Figures

**Supplemental Figure 1:**
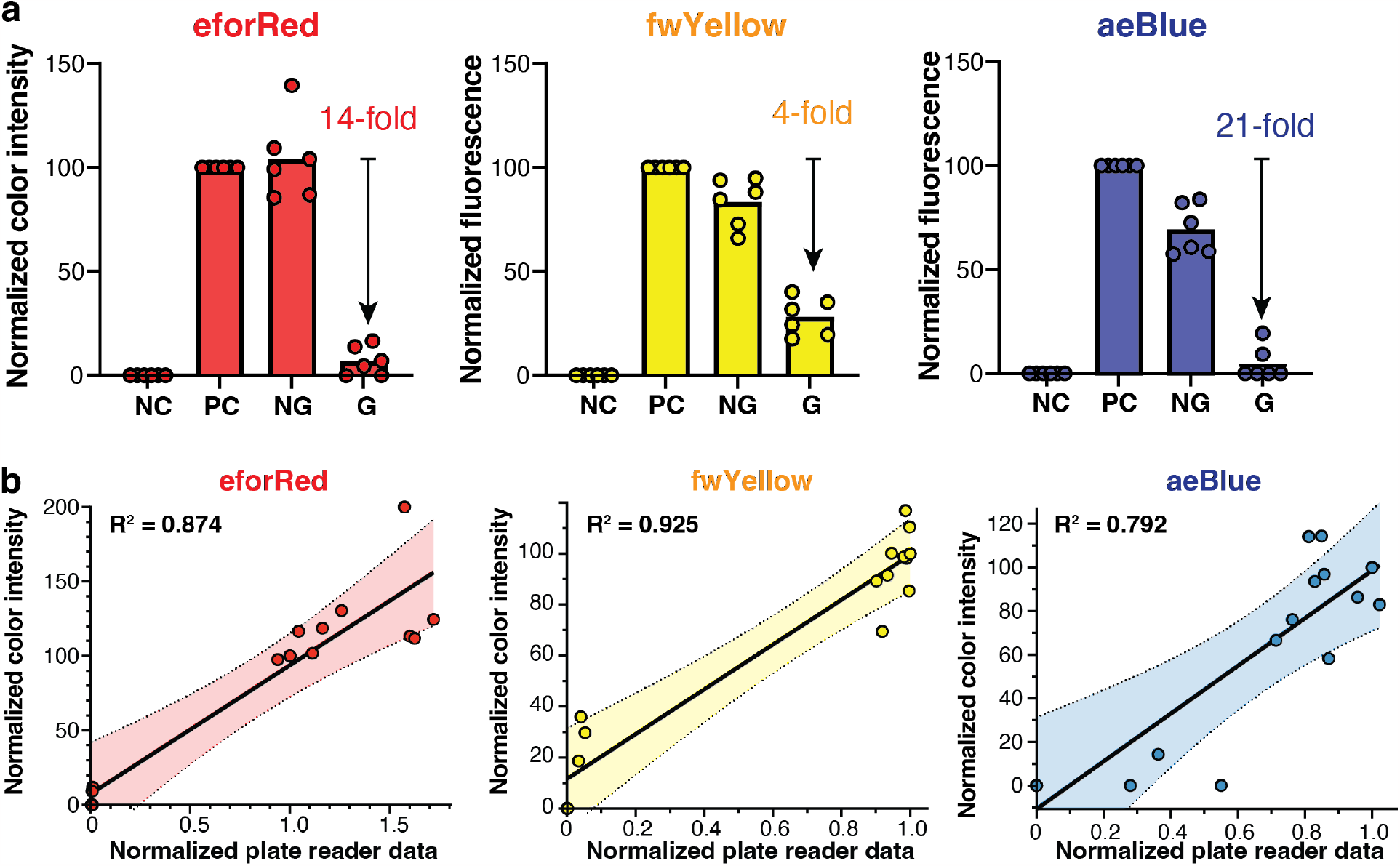
Phone image characterization of the high-tech single-color CRISPR kit reactions. Related to Figure 2d. **a**. Experimental data showing the performance of high-tech single-color CRISPR kit reactions measured by smartphone imaging. For each chromoprotein, we compare four conditions: negative control (NC) for water only, positive control (PC) containing CFS and the chromoprotein plasmid, no sgRNA condition (NG) containing CFS, the chromoprotein plasmid, dCas9 protein and an empty sgRNA plasmid, and targeting sgRNA condition (G) containing CFS, the chromoprotein plasmid, dCas9 protein and the cognate sgRNA. For each condition, three biological replicates with readings from the fluorescence plate reader are shown. The bars represent the mean. **b**. 2D scatter plots showing comparison of fluorescence plate reader data (x-axis) and phone image data (y-axis) for eforRed, fwYellow, and aeBlue. Individual dots represent biological replicates in each condition as shown in **Fig. 2d** and **Fig. S1a**. Black lines represent the linear fit and the dotted lines represent 99.9% confidence interval. R^2^ values are shown in the diagrams for the linear regression.

**Supplemental Figure 2:**
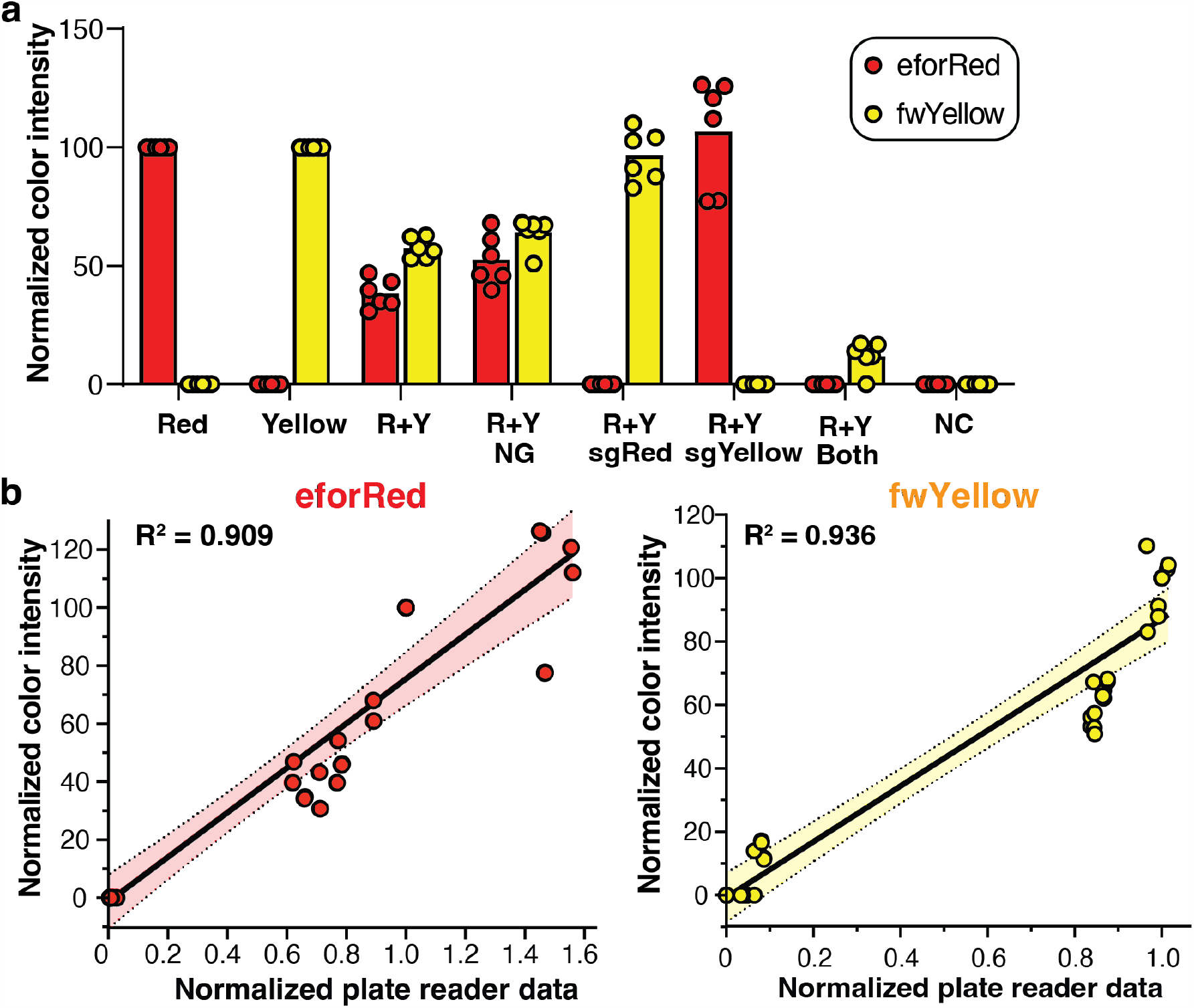
Phone image characterization of the high-tech dual-color CRISPR kit reactions. Related to Figure 3b. **a**. Experimental data showing the performance of high-tech dual-color CRISPR kit reactions measured by smartphone imaging. From left to right, each group of bars (red – eforRed, yellow – fwYellow) show eforRed single-color positive control, fwYellow single-color positive control, eforRed + fwYellow dual-color positive control, dual-color with an empty sgRNA plasmid (NG), dual-color with eforRed-targeting sgRNA (sgRed), dual-color with fwYellow-targeting sgRNA (sgYellow), dual-color with both eforRed-and fwYellow-targeting sgRNAs, and negative control (water only). For each condition, three biological replicates and six technical replicates (each biological replicates measured twice using the smartphone) are shown. The bars represent the mean. **b**. 2D scatter plots showing comparison of fluorescence plate reader data (x-axis) and phone image data (y-axis) for eforRed (left) and fwYellow (right). Individual dots represent biological replicates in each condition as shown in **Fig. 3b** and **Fig. S2a**. Black lines represent the linear fit and the dotted lines represent 99.9% confidence interval. R^2^ values are shown in the diagrams for the linear regression.

**Supplemental Figure 3:**
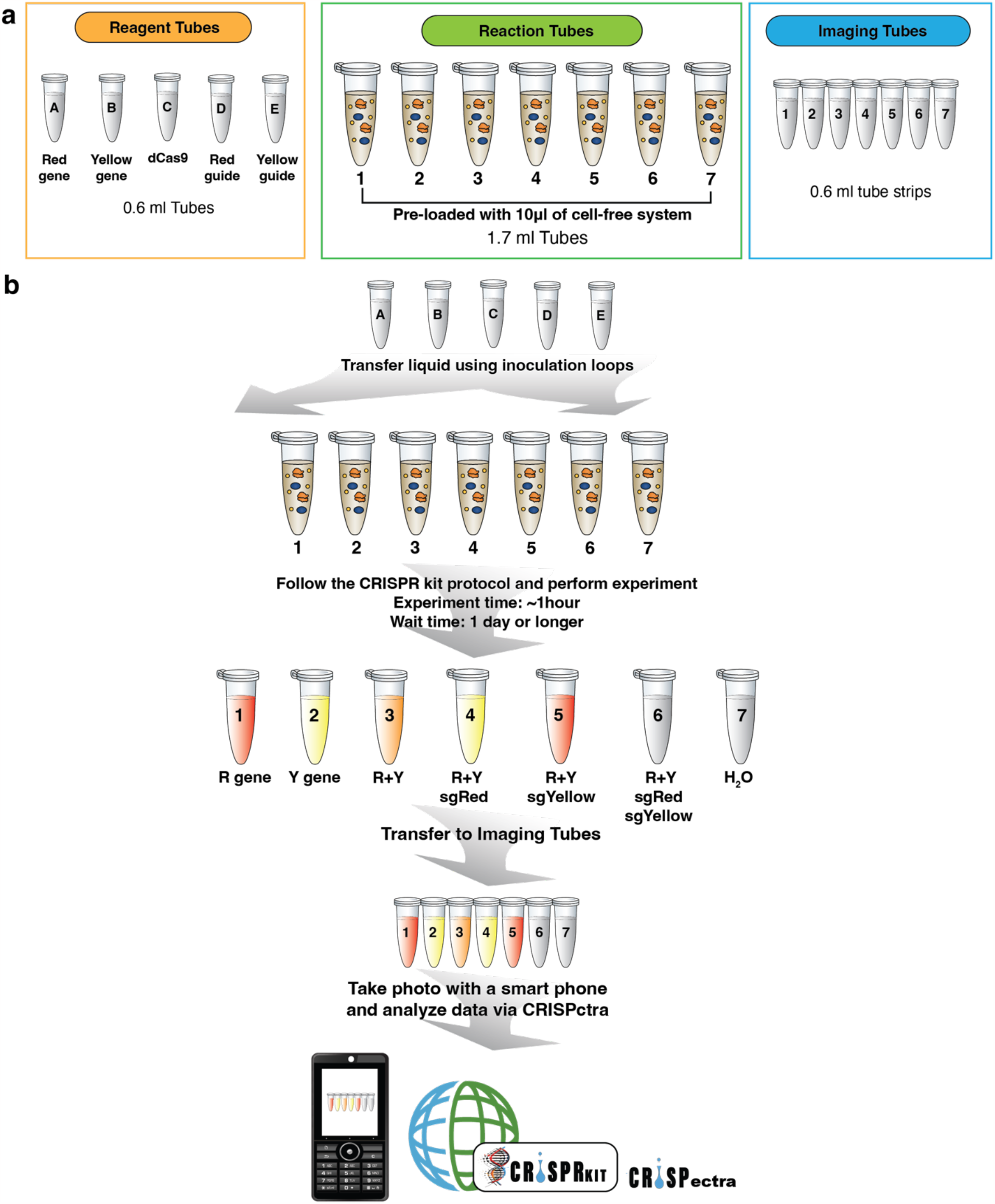
The components and procedure of the frugal dual-color CRISPR kit experiment. Related to Fig. 5a. **a**. Illustration of reagent tubes, reaction tubes, and imaging tubes. **b**. Illustration of the experimental procedure for the protocol for performing dual-color CRISPR kit reactions.

**Supplemental Figure 4:**
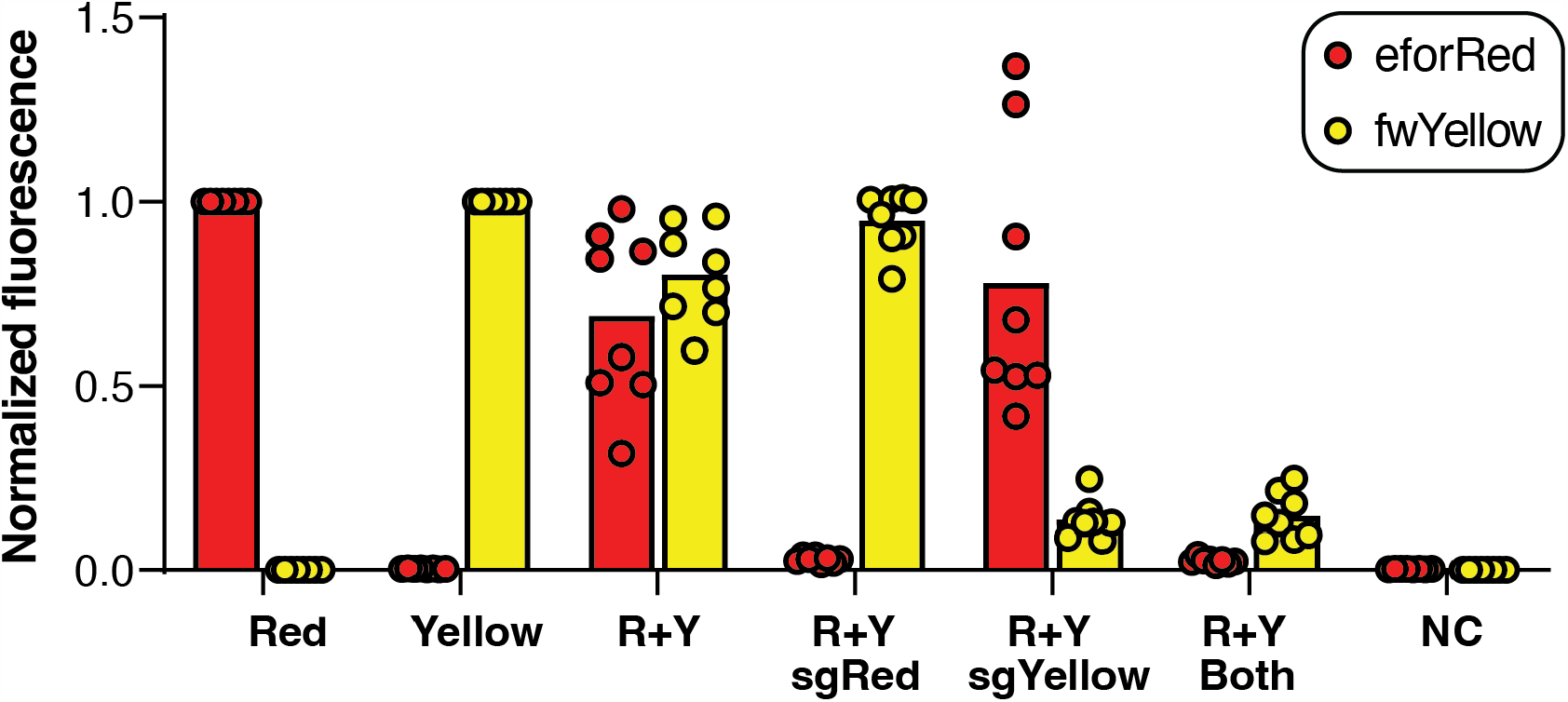
Fluorescence plate reader measurement of the frugal dual-color CRISPR kit reactions performed by high school students. Related to Fig. 5b and 5c. Shown in the diagram is experimental data of the frugal dual-color CRISPR kit reactions measured by a fluorescence plate reader. A total of nine sets of experiments were performed. Eight sets of data were shown, and one experiment failed. From left to right, each group of bars (red – eforRed, yellow – fwYellow) show eforRed single-color positive control, fwYellow single-color positive control, eforRed+fwYellow dual-color positive control, dual-color with the eforRed-targeting sgRNA (sgRed), dual-color with the fwYellow-targeting sgRNA (sgYellow), dual-color with both eforRed- and fwYellow-targeting sgRNAs, and negative control (water only). For each condition, eight experimental replicates (each data point measured twice as technical replicates of phone image analysis) are shown. The bars represent the mean.

**Supplemental Figure 5:**
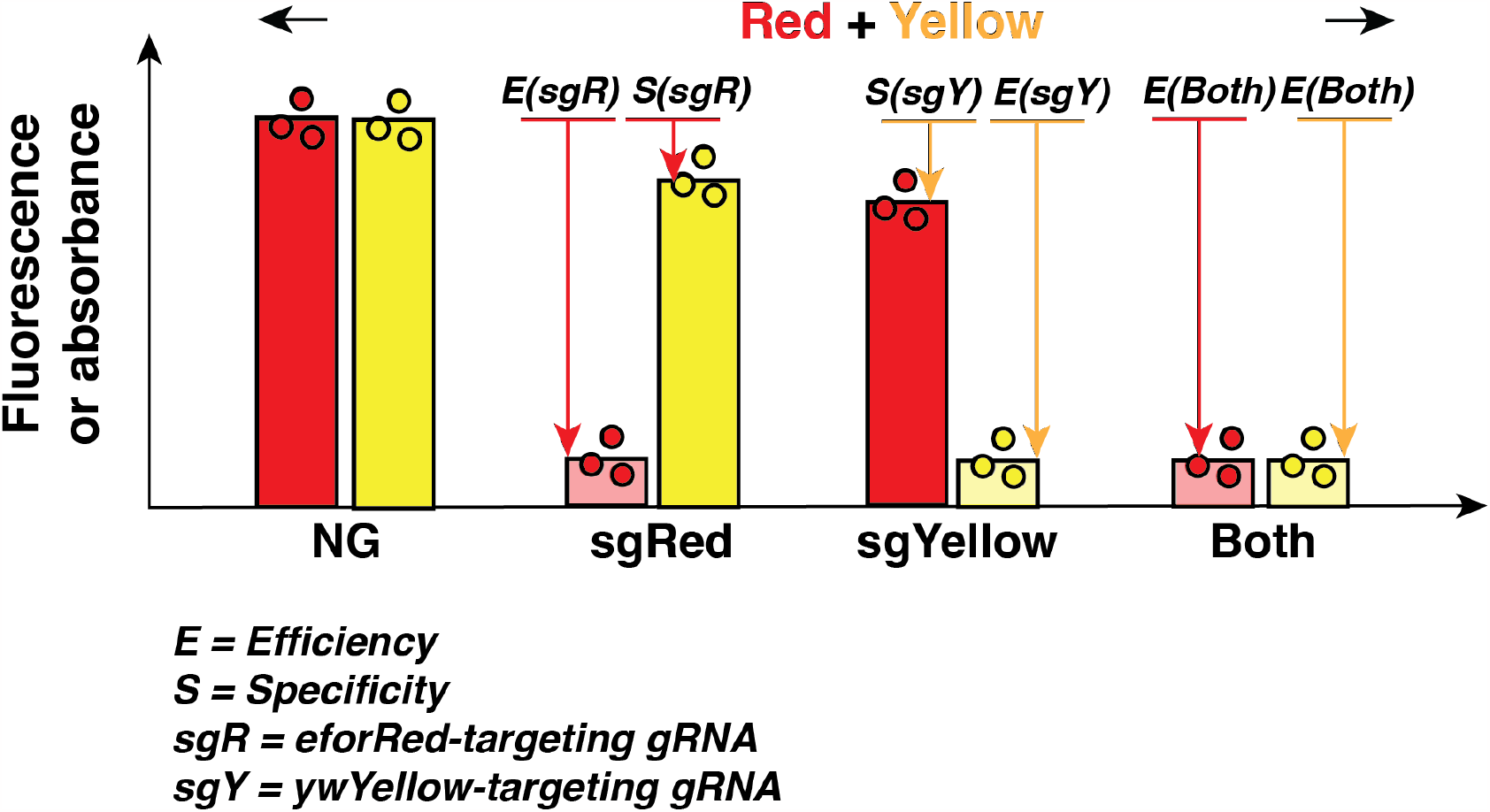
Schematic showing the calculation of efficiency (E) and specificity (S) in dual-color CRISPR kit reactions. Related to Fig. 5d. The diagram uses eforRed and fwYellow as an example. sgR, eforRed-targeting sgRNA; sgY, fwYellow-targeting sgRNA. Both, both eforRed- and fwYellow-targeting sgRNAs.

**Supplemental Figure 6:**
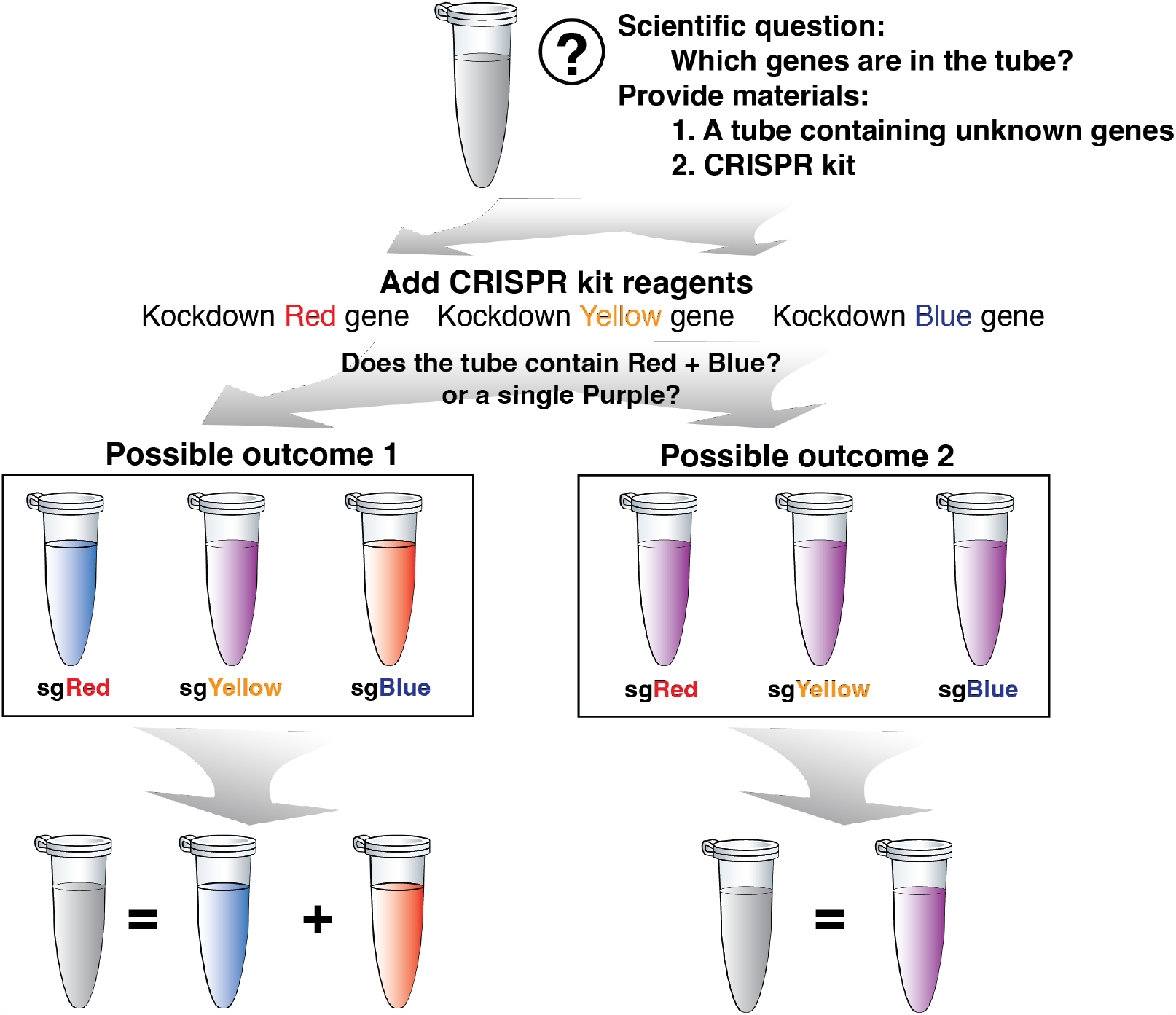
A solution for the ‘guess-and-test’ experiment to figure out the unknown genes. Related to Fig. 6b-6e.

**Supplemental Figure 7:**
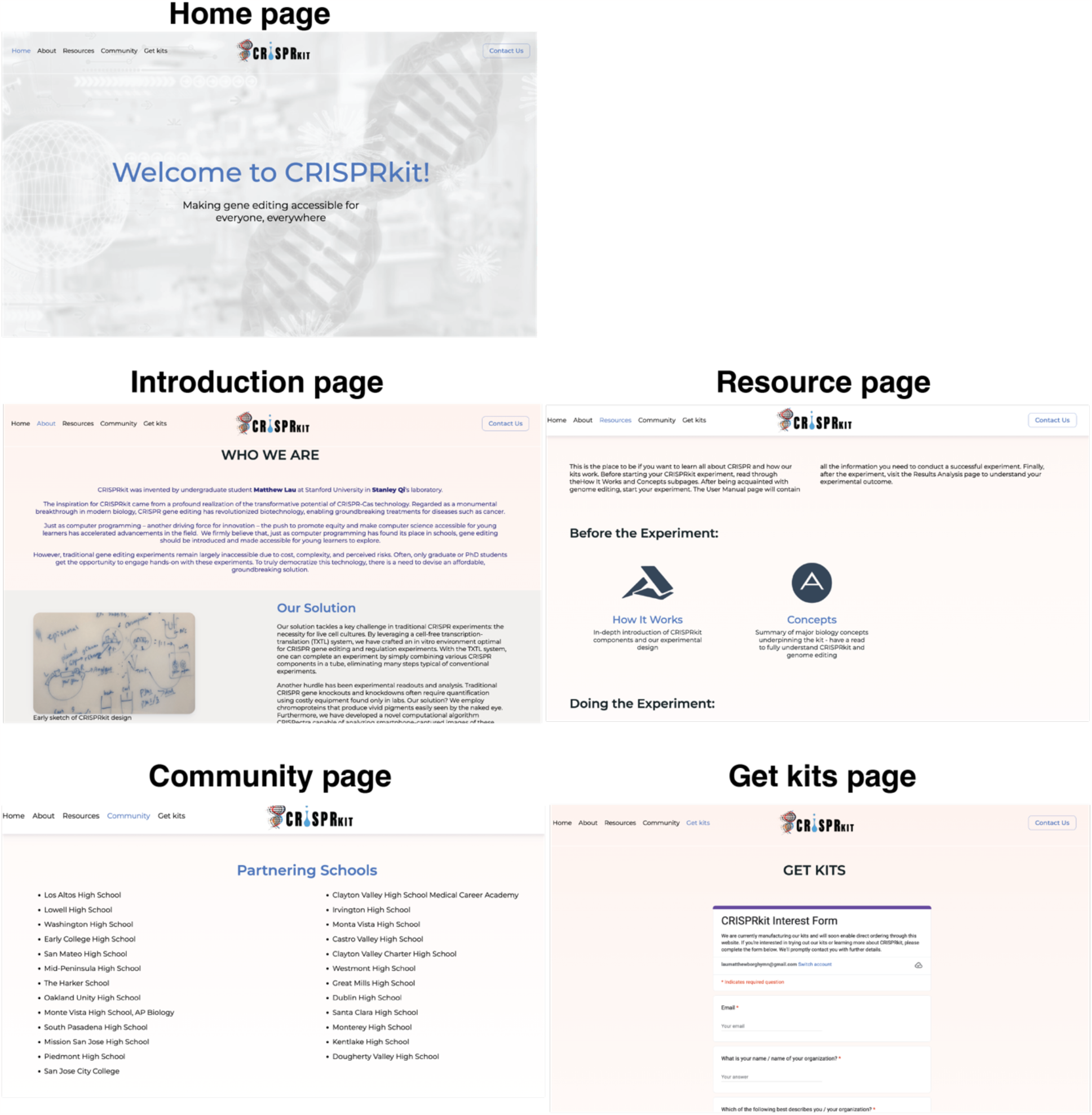
Summary of our CRISPR kit webserver. Multiple screenshots of the website are shown.

## Supplemental Tables

**Table S1:** Plasmids used in the study.

| Plasmid | Sequence Source | Description | Source |
| --- | --- | --- | --- |
| pSLQ3533 | Ref. 22 | J23119 promoter-strong RBS-eforRed-terminator-AmpR-ColE1 | This Work |
| pSLQ3534 | Ref. 23 | J23119 promoter-strong RBS-feYellow-terminator-AmpR-ColE1 | This Work |
| pSLQ14101 | Addgene #117846 | J23119 promoter-strong RBS-aeBlue-terminator-AmpR-ColE1 | This Work |
| pSLQ3538 | --- | J23119 promoter-eforRed-targeting sgRNA-terminator-AmpR-ColE1 | This Work |
| pSLQ14103 | --- | J23119 promoter-aeBlue-targeting sgRNA-terminator-AmpR-ColE1 | This Work |
| pSLQ14107 | --- | J23119 promoter-fwYellow-targeting sgRNA-terminator-AmpR-ColE1 | This Work |
| pSLQ230 | Ref. 18 | J23119 promoter-terminator-J23119 promoter-terminator-AmpR-ColE1 | This Work |

**Table S2:**
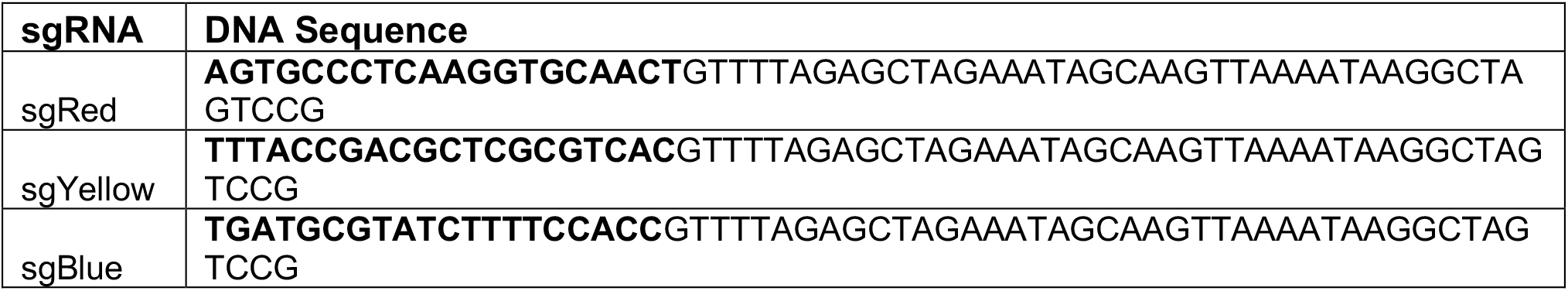
Sequences of sgRNAs. Bold region indicates the DNA-targeting sequence.

**Table S3:**
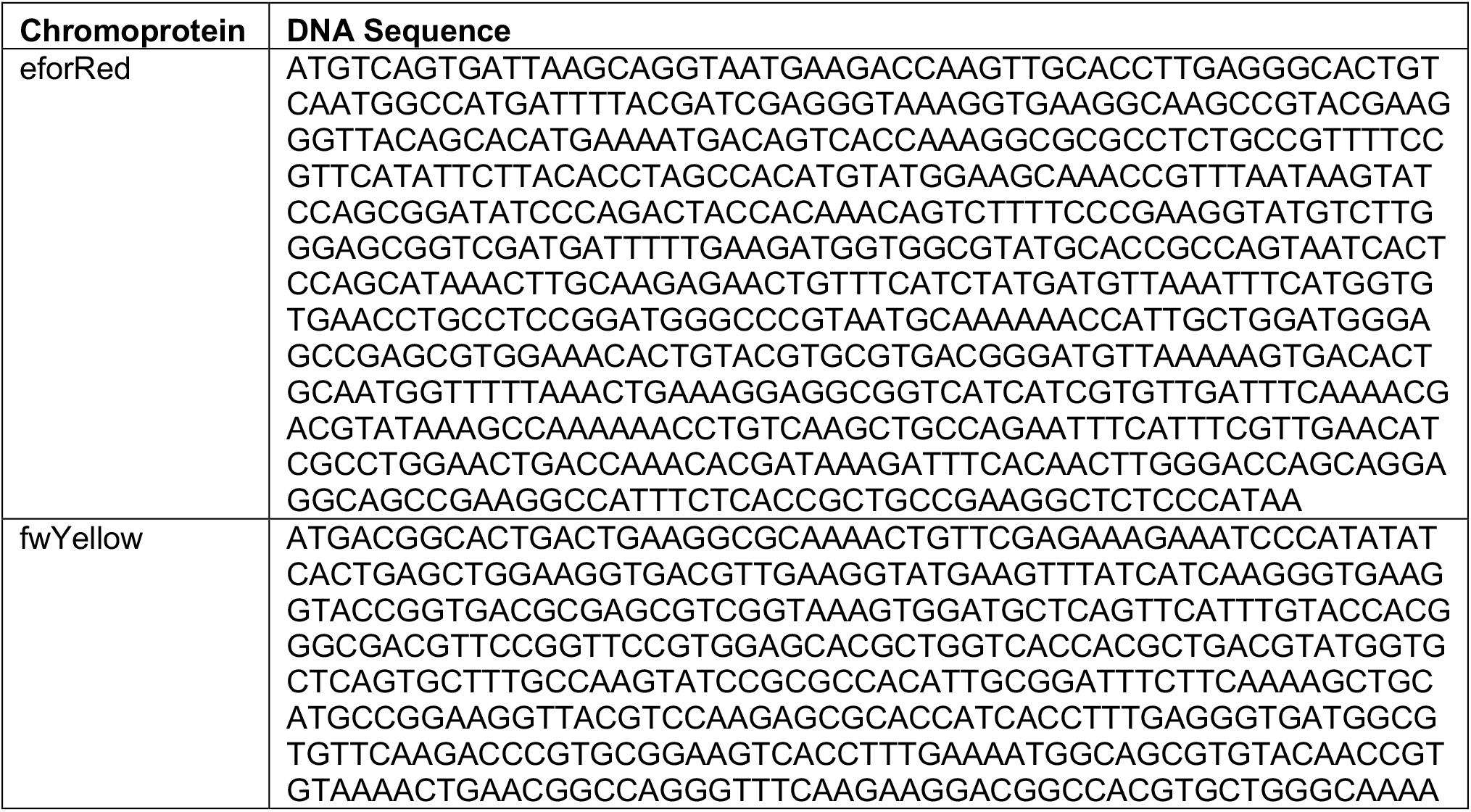
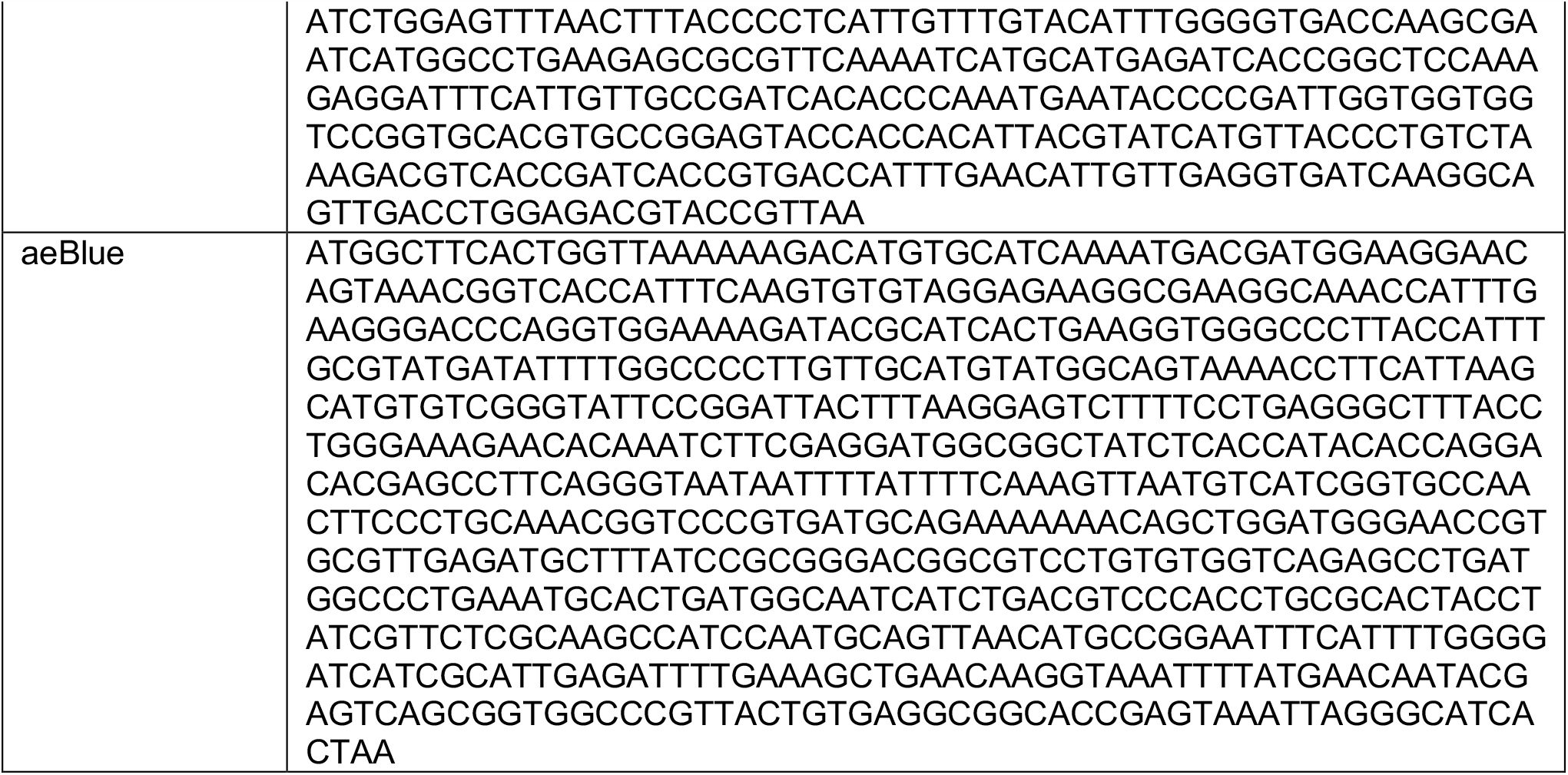
DNA sequences of chromoproteins.

**Table S4:**
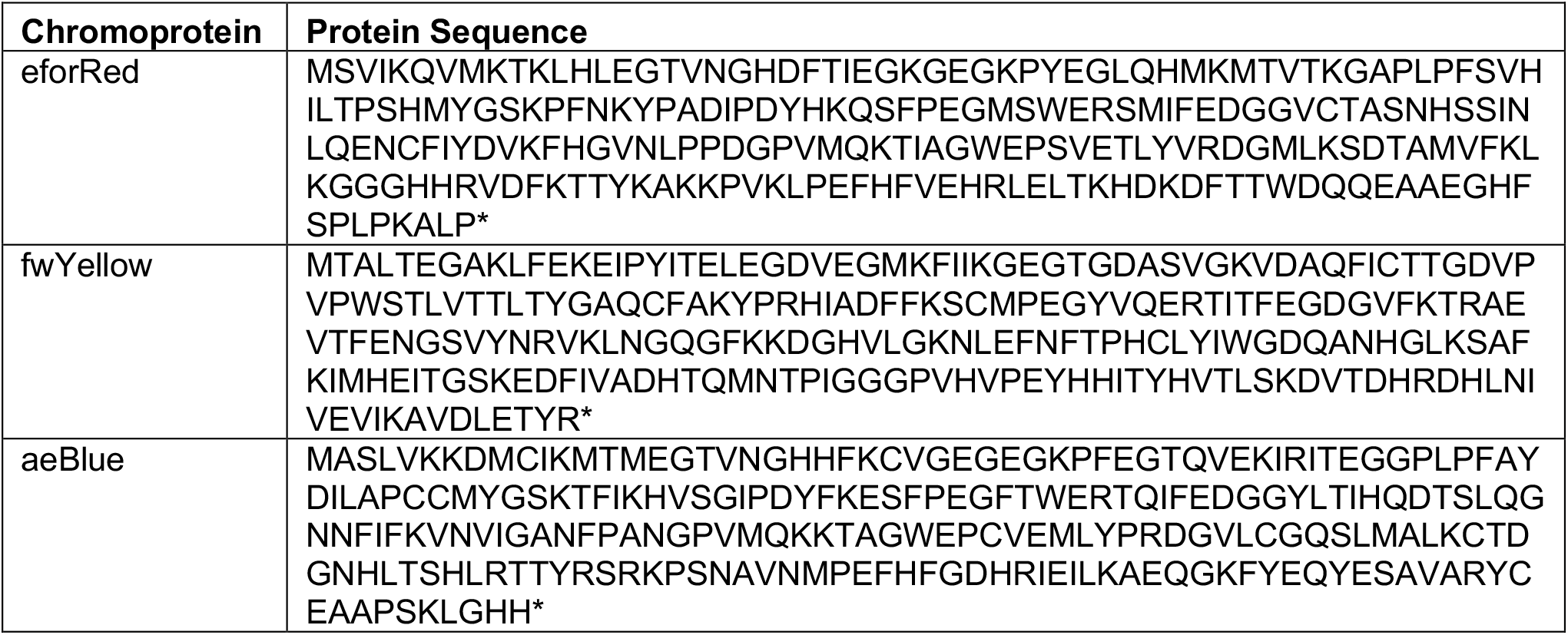
Protein sequences of chromoproteins.

## Supplemental Movies

Movie S1: Tutorial movie for performing the CRISPR kit experiment

Movie S2: Tutorial movie for analyzing the CRISPR kit experiment via smartphone

## Supplemental Notes

### Supplemental Note 1: Core concepts related to the CRISPR kit

These are the essential concepts that form the knowledge basis of using the CRISPR kit. Delve into them to understand the workings of the kit and its underlying processes, especially if these concepts are relatively new to you.

### Biological concepts

#### DNA

DNA, Deoxyribonucleic Acid, is the blueprint of life, holding the genetic instructions that guide growth, development, function, and reproduction of living organisms. DNA’s structure, a double helix, looks like a twisted ladder where the rungs are composed of pairs of nucleotides - adenine (A) with thymine (T), and cytosine (C) with guanine (G).

#### RNA

RNA, Ribonucleic Acid, much like DNA, is a chain of nucleotides playing a critical role in translating the genetic information from DNA into proteins - life’s building blocks. RNA, though, is single-stranded, are composed of nucleotide uracil (U) instead of thymine (T) found in DNA. Messenger RNA (mRNA), Transfer RNA (tRNA), and Ribosomal RNA (rRNA) are the three primary types of RNA working in unison to create proteins.

#### Protein

Proteins, the fundamental building blocks of life, are large, complex molecules vital for the structure, function, and regulation of the body’s tissues and organs. Formed by linking amino acids in a sequence determined by DNA, proteins perform a myriad of functions from providing structural support to acting as enzymes, facilitating transport, enabling cell signaling and communication, and aiding in defense against foreign substances.

In our kit, we use a type of chromoprotein, which produces a protein that contains a pigment. Different chromoproteins have been characterized from Nature with various colors.

#### Chromoprotein

A chromoprotein is a protein that contains a colored chromophore group, which absorbs particular wavelengths of light, resulting in the protein exhibiting a specific color. These chromophores can be either covalently attached or non-covalently bound to the protein.

#### The Central Dogma

The Central Dogma outlines the fundamental flow of genetic information within an organism, comprising:

1. DNA replication: Cells create an exact copy of their DNA prior to division.
2. Transcription: DNA’s genetic instructions are transcribed into messenger RNA (mRNA).
3. Translation: mRNA guides the assembly of proteins in the cytoplasm.

This sequential process, from DNA replication to protein synthesis, is essential to life, driving the functionality of an organism’s cells.

#### Plasmid

A plasmid is a small, circular, double-stranded DNA molecule that is distinct from the chromosomal DNA within a cell. Plasmids are commonly found in bacteria, but they can also be observed in archaea and some eukaryotic organisms.

#### Gene

A gene is a segment of DNA, housing specific sequences of nucleotide pairs that dictate the formation of proteins. These proteins are cellular workhorses, tasked with diverse functions like structure provision, regulation of processes, and facilitation of chemical reactions.

#### Gene Expression

If we regard gene is a language of life, gene expression is how this language is spoken. A language that cannot be spoken is meaningless to life.

Gene expression is the process through which the instructions within our DNA are used to create proteins. Gene expression is tightly regulated process, which ensures the appropriate genes are active at the right time, in the right cells. A standard gene expression process involves two steps: transcription and translation.

### Technological concepts

#### Genome engineering

Genome engineering is a powerful technique that enables precise control or modification to an organism’s DNA. If we regard DNA as the blueprint text of life, genome engineering is the process to alter the text and its meaning. Genome engineering has powerful applications in numerous fields, including disease treatment, agriculture, manufacture of valuable organisms and compounds, environment, climate, and ecology.

#### CRISPR

CRISPR, or “Clustered Regularly Interspaced Short Palindromic Repeats,” is a naturally occurring bacterial defense system against viruses, repurposed by scientists for gene editing.

This system comprises two components: Cas9 protein and single guide RNA (sgRNA). The sgRNA, akin to a mailman, precisely directs Cas9 to the specific DNA segment targeted for modification.

#### CRISPR-Cas9 genome editing

As an important and specific branch of genome engineering technologies, genome editing refers to sequence modifications of the DNA, much like using molecular scissors.

Inside a cell, Cas9 and gRNA collaborate to locate and cut the target DNA sequence, triggering the cell’s inherent repair mechanisms. Two main repair pathways can be leveraged: non-homologous end joining (NHEJ) and homology-directed repair (HDR). NHEJ can cause small sequence alterations, ideal for gene disruption or inactivation, while HDR, given a DNA template, facilitates precise DNA sequence alterations or insertions at the targeted site. CRISPR genome editing represents a powerful tool for advancing biological understanding and potential health benefits.

#### CRISPR-dCas9 genome regulation

CRISPR-dCas9 uses a modified Cas9 protein, “dead” Cas9 (dCas9), to regulate gene expression without altering the DNA sequence. It acts as a switch button on genes, toggling genes on or off. dCas9 can bind to specific DNA sequences guided by a complementary sgRNA but lacks the cutting ability of the original Cas9 protein. This binding prevents the normal genetic machinery from reading and activating the gene, effectively regulating its expression.

To control gene expression, dCas9 can be combined with additional components, such as activators or repressors. For example, we can attach proteins called activators to dCas9 to enhance gene expression, causing the gene to be turned on. Conversely, we can attach repressor proteins to dCas9 to suppress gene expression, turning the gene off. In this kit, we utilize CRISPR-dCas9 gene regulation to suppress expression of various chromoprotein genes.

#### CRISPRi

CRISPRi, or CRISPR interference, is a modified application of the standard CRISPR-Cas system. While the traditional CRISPR-Cas9 approach is used for gene editing by creating double-stranded breaks in the DNA, CRISPRi is designed to modulate gene expression without making permanent changes to the DNA sequence. CRISPRi provides a reversible method to modulate gene expression. This means genes can be “turned off” temporarily without permanently altering their sequences. Besides, CRISPRi offers a high degree of specificity, reducing off-target effects, and has been broadly used for studying gene function, as genes can be silenced to understand their roles in various cellular processes.

#### Cell-free system

A cell-free system refers to an experimental system that studies biological reactions outside of living cells. Instead of using intact cells, the cellular components (like proteins, nucleic acids, and small molecules) are extracted, and the reactions are carried out in a test tube or other suitable containers.

#### Fluorescence quantification

The process of measuring and quantifying the intensity of emitted light (fluorescence) from a sample (e.g., that contains a fluorescence protein or a chromoprotein) after it has been excited with a specific wavelength of light. This method capitalizes on the properties of fluorescent molecules (fluorophores) which, when exposed to a specific excitation wavelength, emit light at a longer wavelength.

### Supplemental Note 2: Protocol for the CRISPR kit

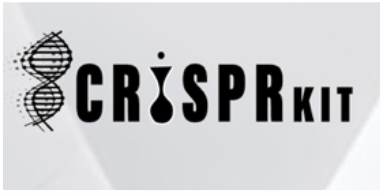

**DAY 1:**

Each kit contains 5 small reagent tubes (A-E), 7 big reaction tubes (1-7), and 7 small imaging tubes (1-7). The reaction tubes are preloaded with the cell-free system (1-6) or H2O (7). The kit looks like:

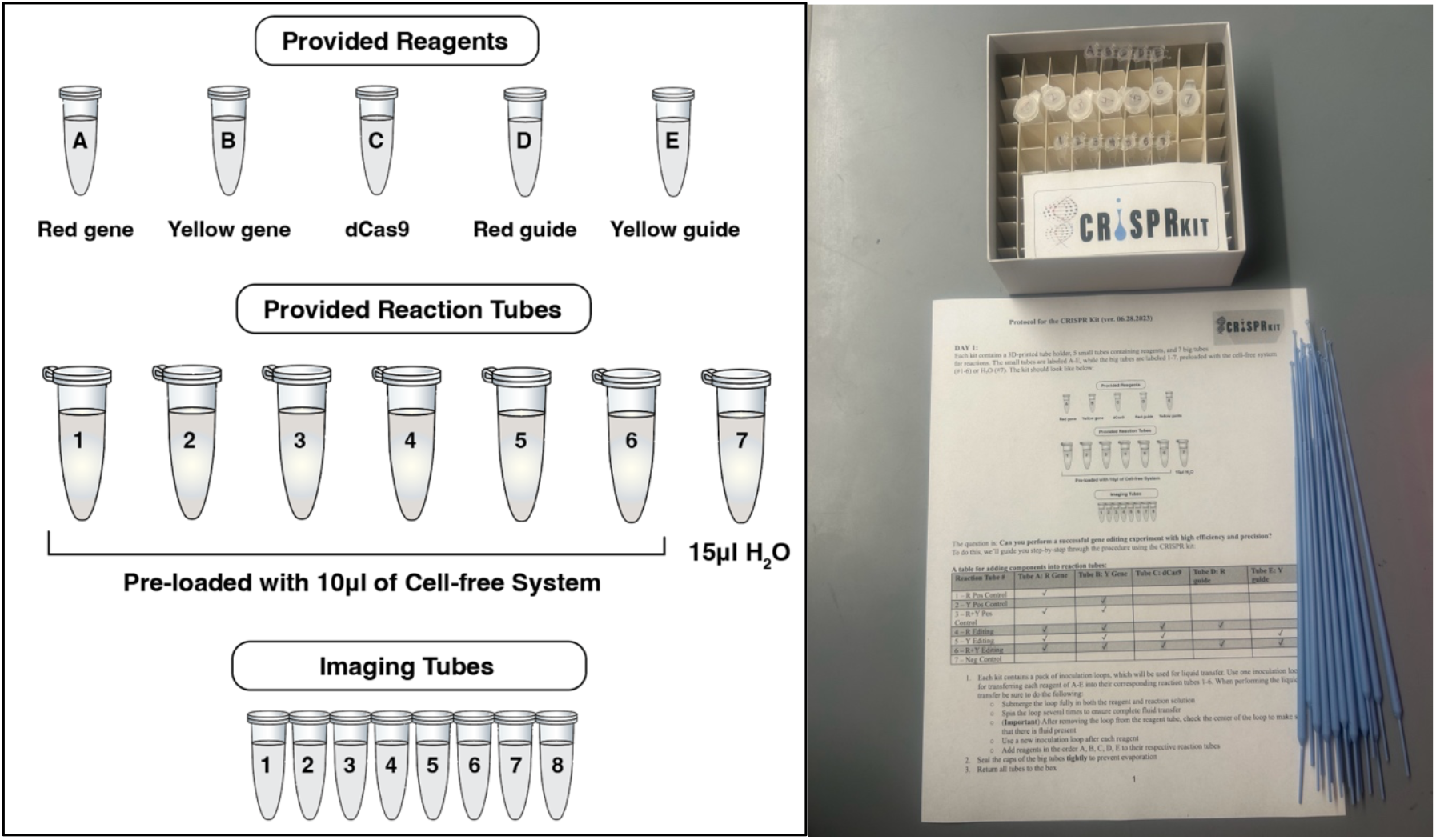

The question is: **Can you perform a CRISPR experiment with high efficiency and precision?**

To do this, we’ll guide you step-by-step through the procedure using the CRISPR kit:

### A table for adding components into reaction tubes

A table for adding components into reaction tubes:
| Reaction Tube # | Tube A:<br>Red Gene | Tube B:<br>Yellow Gene | Tube C:<br>dCas9 | Tube D:<br>Red sgRNA | Tube E:<br>Yellow sgRNA |
| --- | --- | --- | --- | --- | --- |
| 1 – Red Positive Control | ✓ |  |  |  |  |
| 2 – Yellow Positive Control |  | ✓ |  |  |  |
| 3 – R+Y Positive Control | ✓ | ✓ |  |  |  |
| 4 – R+Y+sgRed | ✓ | ✓ | ✓ | ✓ |  |
| 5 – R+Y+sgYellow | ✓ | ✓ | ✓ |  | ✓ |
| 6 – R+Y+sgRed+sgYellow | ✓ | ✓ | ✓ | ✓ | ✓ |
| 7 – Negative Control (H<sub>2</sub>O) |  |  |  |  |  |

1. Each kit contains a pack of inoculation loops (20), which will be used for liquid transfer. Use one inoculation loop for transferring each reagent of A-E into their reaction tubes:
  a. Submerge the loop fully in both the reagent and reaction solution
  b. Spin the loop several times to ensure complete fluid transfer
  c. (**Important**) After removing the loop from the reagent tube, check the center of the loop to make sure that there is fluid present
  d. Use a new inoculation loop after each liquid transfer
  e. Add reagents in the order A, B, C, D, E to their respective reaction tubes
2. Seal the caps of the big tubes **tightly** to prevent evaporation, and return all tubes to the box
3. Allow the reactions to proceed for more than 16 hours. Any time between 16-48 hours is fine

**Day 2:** On Day 2, you will analyze the results of the reactions. Your reactions should look like this (Explain to your teacher why):

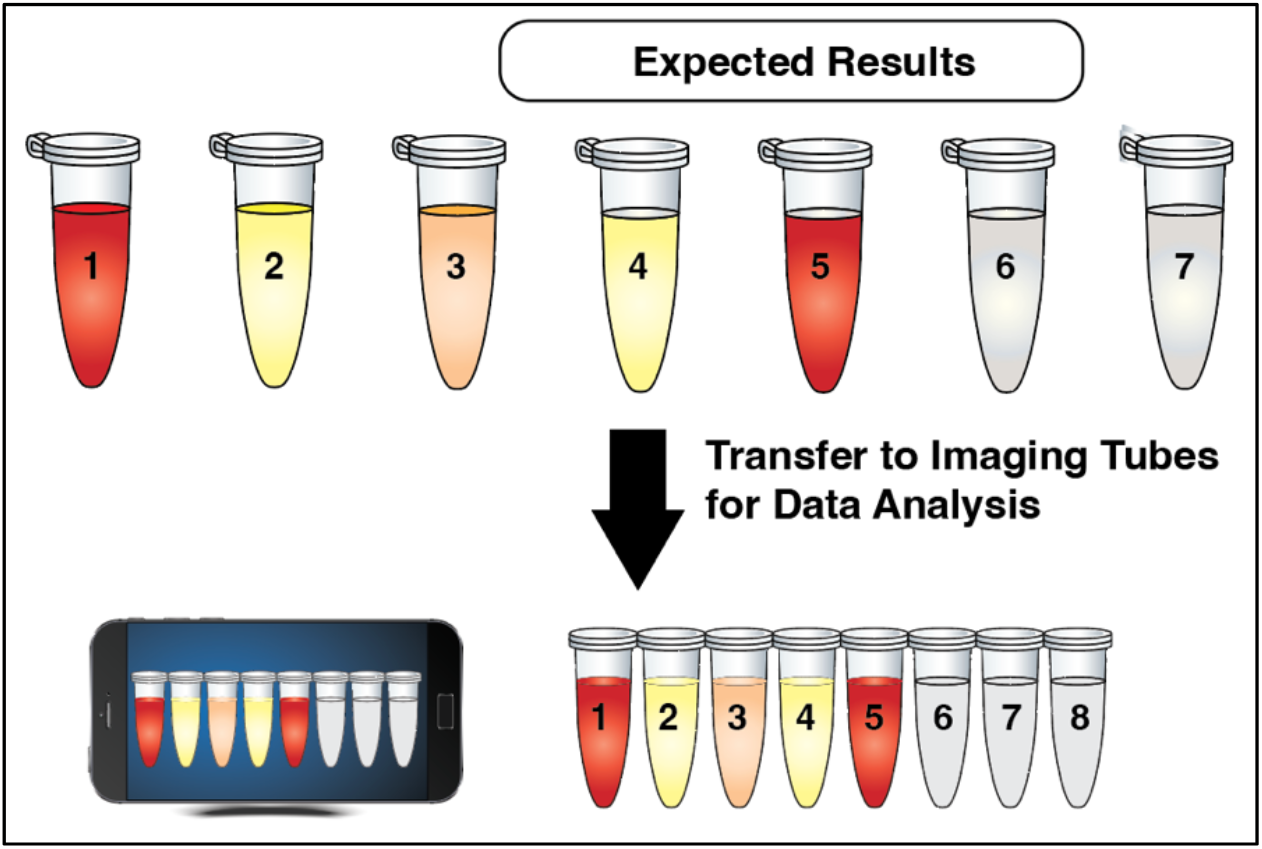

1. For better imaging analysis, transfer all reaction in each tube to the corresponding Imaging Tubes (small).
2. Carefully lay the reaction tubes on a piece of white paper. Take a picture with flash on. Airdrop/send it to your laptop. (**Important**) be sure to minimize glare and shadows! ***Example***:

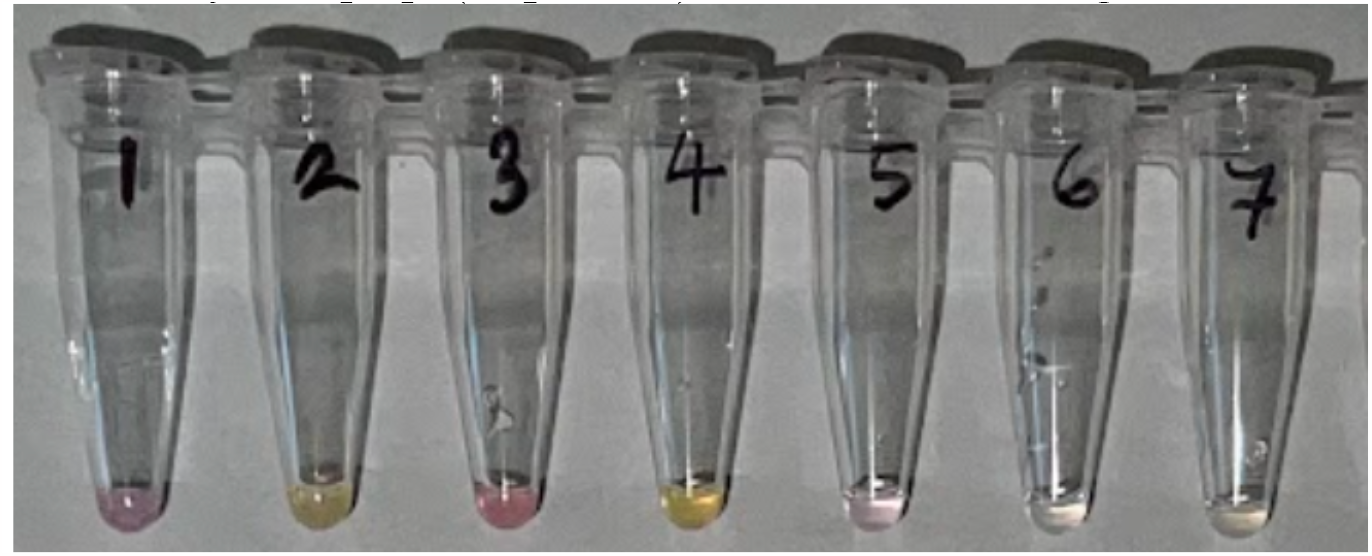
3. Use the CRISPectra algorithm for image analysis.

## Appendix 1 1: Contents of the CRISPR kit

1. dCas9 protein
2. Chromoprotein plasmids encoding Red and Yellow
3. Two sgRNA plasmids each targeting Red and Yellow
4. 5x Reagent Tubes (small)
5. 7x Reaction Tubes (big), preloaded with the cell-free system
6. 7x Imaging Tubes (small)
7. Manual and educational content

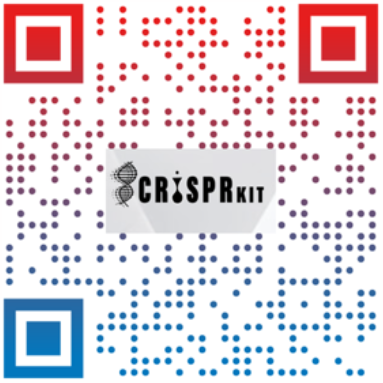

## Appendix 2 The CRISPR kit website: http://crisprkit.org

